# Sorting motifs target the movement protein of ourmia melon virus to the trans-Golgi network and plasmodesmata

**DOI:** 10.1101/724716

**Authors:** Natali Ozber, Paolo Margaria, Charles T. Anderson, Massimo Turina, Cristina Rosa

## Abstract

Plants have a highly sophisticated endomembrane system targeted by plant viruses for cell-to-cell movement. The movement protein (MP) of ourmia melon virus (OuMV) is delivered to plasmodesmata (PD) and forms tubules to facilitate cell-to-cell movement. Although several functionally important regions for correct subcellular localization of OuMV MP have been identified, little is known about the pathways OuMV MP hijacks to reach the PD. Here, we demonstrate that OuMV MP localizes to the trans-Golgi network (TGN), but not to the multivesicular body/prevacuolar compartment or Golgi, and carries two putative sorting motifs, a tyrosine (Y) and a dileucine (LL) motif, near its N-terminus. Glycine substitutions in these motifs result in loss of OuMV infectivity in *Nicotiana benthamiana* and Arabidopsis. Live cell imaging of GFP-labeled sorting motif mutants shows that Y motif mutants fail to localize to the TGN, plasma membrane, and PD. Mutations in the LL motif do not impair plasma membrane targeting of MP but affect its ability to associate with callose deposits at the PD. Taken together, these results suggest that both motifs are indispensable for targeting OuMV MP to PD and for efficient systemic infection but show differences in functionality. Co-immunoprecipitation assays coupled with mass spectrometry identified a series of host factors that could interact with the OuMV MP and link the MP with various pathways, in particular vesicle trafficking and membrane lipids. This study provides new insights into the intracellular targeting of MPs and pathways that plant viruses hijack for cell-to-cell movement.

## Introduction

Viral movement proteins (MPs) are critical for mediating virus movement through plasmodesmata (PD) as the size of virus nucleic acids and virions does not allow for passive intercellular movement. Indeed, localization of viral MPs to PD has been demonstrated, and some MPs can increase the PD size exclusion limit to enable the movement of viruses into adjacent cells (1,2). Viral MPs use various transport strategies to reach PD and interact with several host factors to regulate movement processes (3–5). A well-studied viral MP, tobacco mosaic virus (TMV) MP, recruits cytoskeleton components (6–9), host factors involved in the intra- and intercellular transport (10–12), and cellular membranes (13). Unlike TMV MP, substantially less host components have been reported for tubule-forming MPs [e.g., grapevine fanleaf virus (GFLV), cauliflower mosaic virus (CaMV), and cowpea mosaic virus (CPMV)] (14–20). Many functionally and structurally diverse proviral host factors, particularly in the plant endomembrane system, remain to be elucidated.

The plant endomembrane system has unique players controlling trafficking pathways to modulate plant-specific cellular functions and stress responses (21,22). The trans-Golgi network (TGN) acts as an early endosome (EE) where the major sorting processes (i.e., secretion, endocytosis, recycling, and vacuolar targeting) are orchestrated (23). Endocytosed cargoes are received by the TGN, and these cargoes are either returned to the plasma membrane or sent to multivesicular bodies (MVBs)/prevacuolar compartments (PVCs), which in plant cells serve as late endosomes (LEs). Heterotetrameric adaptor protein (AP) complexes, which consist of two large subunits (γ/α/δ/ε/ζ and β), a medium subunit (μ) and a small subunit (σ), are key components of protein sorting in endocytic and post-Golgi secretory pathways (24). AP-dependent protein sorting is mediated most commonly by the recognition of sorting signals, tyrosine (Y), YXXØ and dileucine (LL), [D/E]XXXL[L/I] motifs (where Ø is a bulky hydrophobic residue and X any amino acid) on cargo proteins. In plants, one of the most studied sorting signals is the Y motif. Several cell surface receptors, receptor-like kinases, and receptor-like proteins (25), transporters (26), and vacuolar sorting receptors (27–29) carry Y motifs. There are, however, only a few reports showing the recognition of these signals by AP complexes (27– 32). LL motifs in plants are far less studied.

MPs of several viruses contain putative Y and LL motifs. However, the role of these sorting motifs in the targeting of viral MPs to PD is still largely unknown. The GFLV MP contains putative Y and LL motifs, which are also found in MPs of several other nepoviruses (14). The conserved YQDLN motif, which conforms to the Y motif, of the triple gene block protein TGB3 of poa semilatent virus (PSLV) is involved in targeting of the protein to the cell periphery (33). Potato mop-top virus (PMTV) TGB3 also carries the same motif, and the disruption of this motif by introducing mutations impairs the localization of PMTV TGB3 to the ER and motile granules, and the targeting of PMTV TGB3 to PD (34,35). Moreover, three functional Y motifs (i.e., YLPL, YGKF, and YPKF) of CaMV MP govern the endosomal localization of CaMV MP and thus its peripheral presence (16).

The MP of ourmia melon virus (OuMV), a member of the genus *Ourmiavirus*, forms tubules across the cell wall of epidermal cells of OuMV-infected *Nicotiana benthamiana*, harboring assembled virions (36). Confocal microscopy of green fluorescent protein-tagged OuMV MP (GFP:MPwt) also shows the presence of protruding tubular/punctate structures in *N. benthamiana* and *A. thaliana* (37), as well as the presence of GFP:MP puncta between neighboring cells, co-localizing with callose deposits at PD (37,38). Alanine-scanning mutagenesis of conserved amino acids of MPs belonging to the 30KDa family revealed three residues at positions 98, 150, and 169 that are required for targeting of OuMV MP and tubule formation in protoplasts and important for MP structure (37). It is, however, still unknown which intracellular pathways OuMV MP uses to facilitate cell-to-cell movement of virions. OuMV MP contains two potential internalization and trafficking motifs, one putative Y motif, ^88^YDKV, and one putative LL motif, ^59^DPIALI. In this study, we investigated the involvement of OuMV MP in post-Golgi trafficking pathways and characterized the individual function of each motif in relation to these pathways by introducing glycine substitutions into critical residues of these motifs and examining localization of mutants by confocal microscopy. Moreover, we identified host factors potentially recruited by OuMV MP to reach the cell periphery.

## Results

### 1. OuMV MP is associated with vesicle trafficking and lipid binding proteins

OuMV MP-interacting proteins were identified by using purified recombinant MBP-MP as bait to isolate interacting proteins from Arabidopsis leaf extracts and subjecting co-immunoprecipitated proteins to LC-MS/MS analysis. After removing non-specific interacting proteins (MBP-binding proteins), a total of 55 unique detergent-soluble and 146 soluble OuMV MP-interacting were discovered. Although ribosomal proteins, elongation factors, and chloroplast-related proteins could be functionally relevant, especially in the context of viral replication, these proteins were further removed from the analysis to focus on the transport of OuMV MP and the pathways it regulates. Among these candidates, 12 ‘detergent-soluble’ proteins (Table S1) and 35 ‘soluble’ proteins (Table S2) were selected as candidate interacting partners of OuMV MP, representing diverse pathways that OuMV MP could potentially target, including membrane trafficking, calcium signaling, transport across the plasma membrane, protein folding and homeostasis, proteolysis, and metabolism. Of these, nine host proteins are classified in the vesicle trafficking, including components of AP complexes, and three in the lipid binding, suggesting that the MP could associate with the plant endomembrane system. To further elucidate the membrane components OuMV MP could target in the plant, we screened various phospholipid candidates using a protein lipid overlay assay. OuMV MP exhibited affinity specifically for phosphatidylinositol 3-phosphate (Fig. S1).

### 2. OuMV MP is trafficked through post-Golgi transport pathways

We previously investigated the subcellular localization of a OuMV GFP:MPwt fusion in *N. benthamiana* leaf epidermal cells, and demonstrated the presence of fluorescent foci at the cell surface, co-localized with PD (37,38). Further investigation of the localization of GFP:MPwt, expressed together with the RdRp and CP, revealed that GFP:MPwt was associated with peripheral mobile structures (Movie S1). To examine whether OuMV MP localizes to endosomal compartments, we used an endocytic marker, FM4-64, which has been shown to follow endocytic pathways in plants (39–41). GFP:MPwt-marked puncta appeared to colocalize with FM4-64 in *N. benthamiana* protoplasts (Fig. 1A). To identify the nature of the endosomal compartments in which the MP resides, we transiently co-expressed mCherry-fused organelle marker proteins (42), VTI12 (a TGN marker) (43), and RHA1 (a MVB/PVC marker) (44), with GFP:MPwt along with RdRp and CP in *N. benthamiana* leaf epidermal cells. We also applied two well-known inhibitors of vesicle trafficking, brefeldin A (BFA; an ARF GEF inhibitor) and Wortmannin (Wm; a phosphatidylinositol-3 and -4 kinase inhibitor), to these cells to dissect the pathways through which MP is trafficked.

**Figure 1.**
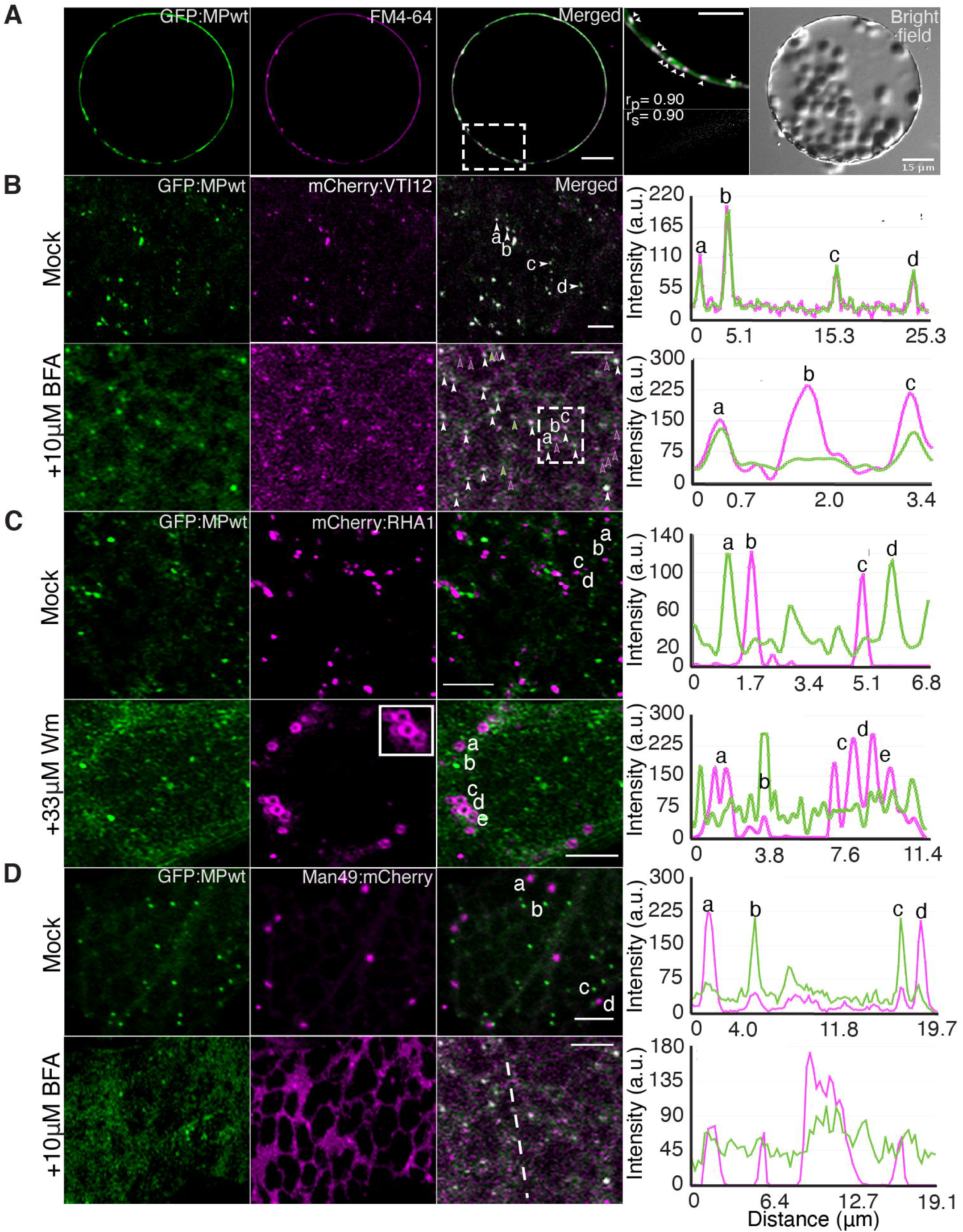
OuMV MP localizes to the TGN/EE, but not to the MVB/PVC or Golgi. **(A)** *N. benthamiana* protoplasts expressing OuMV GFP:MPwt (green) with RdRp and CP. Protoplasts were stained with FM4-64, an endocytic tracer (magenta). A selected region (white rectangle) was analyzed to show the colocalization of MP-labeled punctate structures (arrowheads) with FM4-64. The extent of colocalization was quantified using the PSC plugin in ImageJ. rp, linear Pearson correlation coefficient; rs, nonlinear Spearman’s rank correlation coefficient. Images represent single optical planes. Scale bars, 15 μm, 10 μm (magnified region). **(B-D)** *N. benthamiana* epidermal cells expressing GFP:MPwt (green) with RdRp and CP, and organelle markers (magenta**), (B)** mCherry:VTI12 (TGN/EE), **(C)** mCherry:RHA1 (MVB/PVC) and **(D)** Man49:mCherry (Golgi). Cells were treated with 10 μM BFA (B and D, bottom panel) or 33 μM wortmannin (C, bottom panel) for 1h. White arrowheads indicate colocalized spots, green and magenta arrowheads denote individual GFP:MPwt and mCherry:VTI12 spots, respectively. Images were taken at 66 hpi. Panels represent maximum intensity Z projections. A selected region was analyzed to show the intensity profile generated by ImageJ. Scale bars, 5 μm.

mCherry:VTI12 showed the expected punctate pattern of TGN that overlapped with GFP:MPwt (Fig. 1B), suggesting that MP is recruited to TGN. It has been previously reported that 5-10 μM concentrations of BFA promote dissociation of TGN into small vesicular compartments in tobacco BY-2 cells (45). Hence, we treated *N. benthamiana* leaves expressing both mCherry:VTI12 and GFP:MPwt with 10 µM BFA. Upon BFA treatment, both mCherry:VTI12 and GFP:MPwt dispersed into the cytoplasm. While GFP:MPwt dissociated into smaller punctate structures, some of these structures did not completely co-localize with the TGN marker (Fig 1B, lower panel). When we increased the concentration of BFA to 50 μM, we observed BFA-induced GFP:MPwt aggregates in *N. benthamiana* epidermal cells. We hypothesize that the formation of these aggregates was associated with the toxicity of BFA rather than direct effect of BFA on the trafficking pathways, and we used the lower concentration in all the further experiments.

To assess whether OuMV MP enters a vacuolar trafficking pathway, we investigated the co-localization of GFP:MPwt with mCherry:RHA1. GFP:MP-labeled intracellular compartments did not co-localize with RHA1-positive MVBs/PVCs (Fig. 1C, upper panel). Treatment with 33 μM Wm, which inhibits protein sorting to the vacuole by inducing homotypic fusion and enlargement of MVBs (46,47), resulted in RHA1-positive MVBs/PVCs that appeared as ring-like structures (Fig. 1C, lower panel). These structures were not, however, labeled by GFP:MP (Fig. 1C, lower panel), indicating that MP does not shuttle through the MVB/PVC to the vacuole transport pathway. Unexpectedly, we also observed that the punctate signal of GFP:MP was shifted more toward a diffuse cytoplasmic pattern upon Wm treatment.

Since GFP:MPwt associated with the TGN, we further explored whether the MP puncta were also associated with Golgi bodies by transiently co-expressing GFP:MPwt with Man49:mCherry, a *cis-*Golgi marker (48). As expected, Man49:mCherry displayed a spot-like signal typical of Golgi bodies with a weak signal from the ER; however, the spot-like Golgi signal did not overlap with the punctate signal of GFP:MPwt, indicating that GFP:MPwt structures are distinct from Golgi bodies (Fig. 1D, upper panel) and exclusive to the TGN. We also applied 10 μM BFA to *N. benthamiana* leaves expressing the Golgi marker and GFP:MPwt to further test whether MP sorts in the conventional secretory pathway. In tobacco, BFA inhibits the secretory pathway and causes redistribution of Golgi bodies into ER (49,50). Indeed, treatment with 10 μM BFA caused the Golgi marker to relocate to ER (Fig. 1D, lower panel). While GFP:MP showed a dispersed pattern similar to that observed with the TGN marker upon BFA treatment (Fig. 1B, lower panel), this pattern did not co-localize with the Golgi marker, suggesting that GFP:MPwt does not enter the ER-to-Golgi secretory pathway.

### 3. OuMV MP carries Y and LL motifs, and substitutions into these motifs result in the loss of OuMV infectivity and reduced viral replication

OuMV MP carries one putative Y motif, YDKV (88-91), and one putative LL motif, DPIALI (59–64). To determine whether the Y and LL motifs in OuMV MP are functional, we substituted critical residues of these motifs with glycine and generated five mutants of wild-type MP (MPwt) by site-directed mutagenesis: Y (**G**DK**G**), Y/G (**G**DKV), D (**G**PIA**GG**), D/G (**G**PIALI) and LI/GG (DPIA**GG**), as shown in Fig. 2A. To address possible roles of Y and LL motifs in infectivity of OuMV, the ability of mutants to infect *N. benthamiana* and Arabidopsis (*Arabidopsis thaliana*) was examined using OuMV agroclones carrying the desired mutations. Four out of five OuMV mutants carrying MP_Y, MP_Y/G, MP_D, or MP_LI/GG showed no visible symptoms in local (3 dpi) and systemic (14 dpi) leaves of *N. benthamiana* and Arabidopsis, whereas the OuMV carrying MPwt displayed typical symptoms of leaf chlorosis, curling, and stunted growth (Fig. 2B-C). In Arabidopsis, MP*_*D/G carrying plants exhibited stunted growth as MPwt, but chlorosis and leaf curling were not observed (Fig. 2B, right panel). Unlike in Arabidopsis, mutant MP_D/G occasionally caused mosaic symptoms in systemic leaves of *N. benthamiana*, but the overall symptoms were milder than upon infection with wild-type OuMV (Fig. 2C; Fig. S2). Symptoms of OuMV carrying MPwt or MP_D/G in *N. benthamiana* appeared as early as 5 dpi (Fig. S2) and remained visible during the experimental time frame (14 dpi) (Fig. 2C).

**Figure 2.**
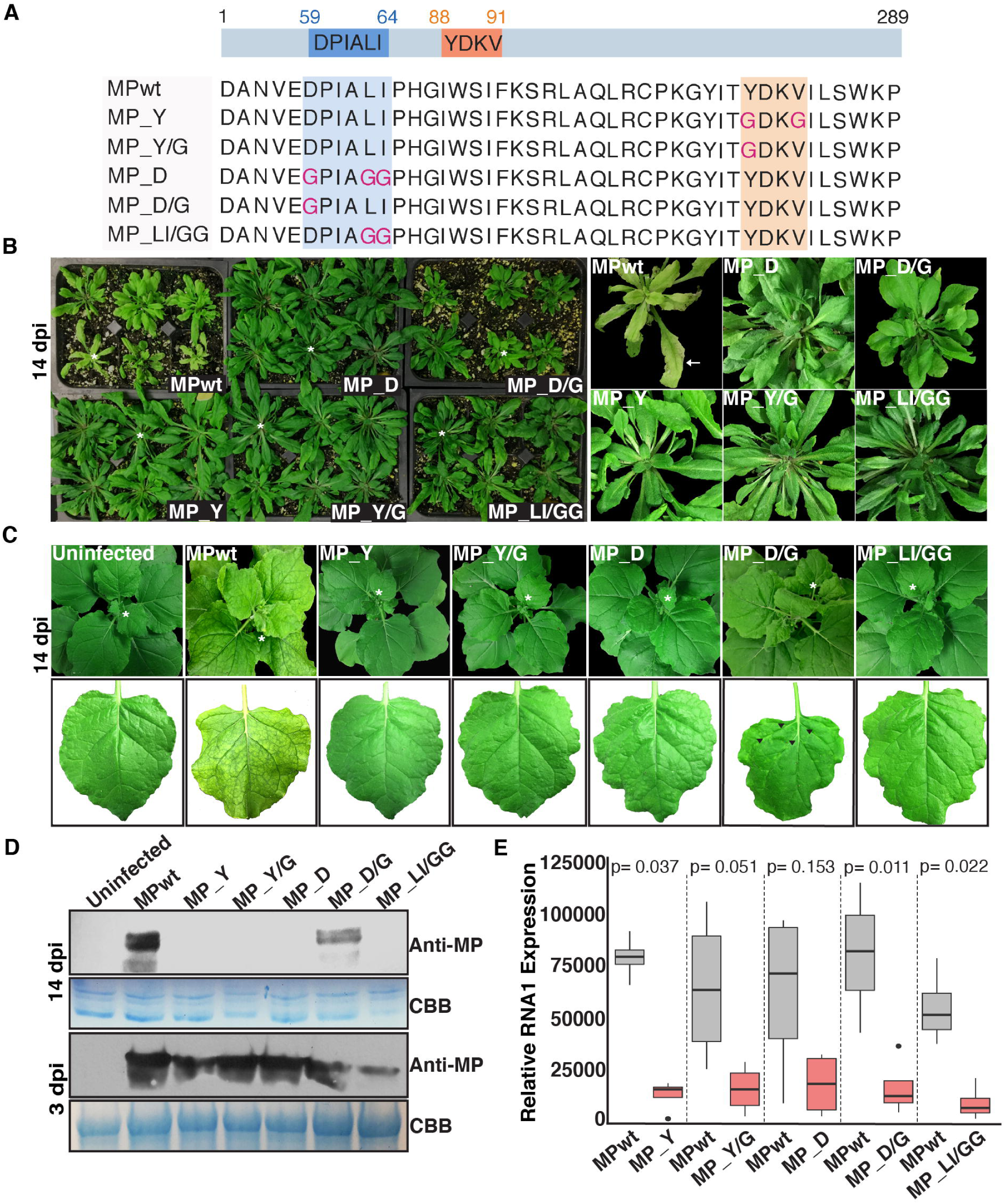
Loss of infectivity and reduced viral replication of OuMV carrying Y and LL motif MP mutants. **(A)** Schematic representation of constructs used in this study. Boxes indicate Y and LL motifs, and mutated sites are shown in pink. The illustration was created with BioRender.com. **(B-C)** Viral symptoms on Arabidopsis **(B)** and *N. benthamiana* **(C)** agroinfiltrated with pGC-RNA1 (RdRp), pGC-RNA3 (CP), and pGC-RNA2 (MP) with indicated mutations, or sterile water (mock) at 14 dpi. White asterisks denote leaves shown in close-up images. Black arrowheads indicate mosaic symptoms on *N. benthamiana* infected with OuMV carrying MP_D/G. The background was removed from the images. **(D)** Western blot analysis of agroinfiltrated leaves at 3 dpi (bottom panel) and upper uninoculated leaves at 14 dpi (top panel). CBB: Coomassie Brilliant Blue, loading control. **(E)** Replication assay of OuMV carrying Y and LL motif MP mutants at 48 hpi. Two leaf spots from 3 leaves were pooled for each sample at 24 hpi and 48 hpi. The qPCR was performed in duplicates. The 2^-ΔΔCt^ method was used to assess fold changes in replication of wild-type and mutants. The data was normalized first using 18S ribosomal RNA, and then a wild-type control at 24 hpi showing the lowest expression. Box plot represents the relative expression of RNA1 (n=4). The statistical analysis was performed on ΔΔCt values using Student’s t-test (unpaired, two-tailed).

The presence of MP mutants in upper uninoculated leaves of *N. benthamiana* at 14 dpi was tested by western blot analysis; only MPwt and MP_D/G could be detected (Fig. 2D, top panel). To test whether the impaired systemic movement of OuMV mutants was due to a lack of expression of MP mutants at the agroinfiltration sites, we performed western blot analysis of *N. benthamiana* leaf extracts for all constructs at 3 dpi, and all MP variants could be detected (Fig. 2D, bottom panel). Virion accumulation was also determined in upper uninoculated leaves at 14 dpi by double antibody sandwich enzyme-linked immunosorbent assay (DAS-ELISA) using α-CP antiserum. All mutants, except the one carrying MP_D/G, failed to move systemically in both *N. benthamiana* and Arabidopsis (Table 1). We also analyzed the coding sequence of MP in OuMV mutants by RT-PCR and sequencing of the PCR product to determine whether mutations in Y and LL motifs were reversed, or whether additional mutations were introduced during infection assays. The reversion of MP_D to wild-type was observed only in one Arabidopsis plant (Table 1), where symptoms became systemic.

**Table 1.** Infectivity assays of OuMV carrying MPwt or MP mutants in *N*.*benthamiana* and Arabidopsis^a^.

| Virus | Systemically infected plants/ total inoculated plants |  |  |  |  |  |
| --- | --- | --- | --- | --- | --- | --- |
|  | *N.benthamiana* |  |  | Arabidopsis (Col-0) |  |  |
|  | Exp. 1 | Exp. 2 | Exp. 3 | Exp.1 | Exp. 2 | Exp. 3 |
| OuMV-MPwt | 6/6 | 6/6 | 6/6 | 8/12 | 9/12 | 7/12 |
| OuMV-MP_Y | 0/6 | 0/6 | 0/6 | 0/12 | 0/12 | 0/12 |
| OuMV-MP_D | 0/6 | 0/6 | 0/6 | 0/12 | 0/11 | 1<sup>*</sup>/12 |
| OuMV-MP_Y/G | 0/6 | 0/6 | 0/6 | 0/12 | 0/12 | 0/12 |
| OuMV-MP_D/G | 6/6 | 5/5 | 6/6 | 6/12 | 8/12 | 6/12 |
| OuMV-MP_LI/GG | 0/6 | 0/6 | 0/6 | 0/12 | 0/12 | 0/12 |

<sup>a</sup>OuMV: *ourmia melon virus*, \*revertant

Next, we assessed if OuMV carrying MP mutants could be inhibited in virus replication. Briefly, we co-infiltrated each MP mutant or MPwt along with RdRp, CP, and pBin61-GFP and collected samples from the infiltrated leaves at 24 hpi and 48 hpi. GFP signal was monitored at 36 hpi to confirm that all plant cells at the infiltrated areas were infected (Fig. S3). Quantification of RNA1 (RdRp) as a proxy for replication was performed by qRT-PCR in all samples. At 24 hpi, RNA1 levels of wild-type OuMV were comparable to that of MP mutants (Fig. S4). While all mutants were able to replicate at high levels at 48 hpi, we observed a substantial decrease in the replication of the mutants compared to corresponding wild-type controls (Fig. 2E). This result suggests that mutations in the OuMV MP Y and LL motifs have also an effect in virus replication.

Given that the μ subunit of Arabidopsis AP2 (AP2M) plays a direct role in the intracellular trafficking of CaMV MP (16), we also investigated the possible role of AP2M in the transport of OuMV MP by infecting Arabidopsis Col-0 (wt), AP2M knockout mutant *ap2m-1* and AP2M-YFP complementation plants with OuMV by mechanical inoculation. There were no obvious differences in symptoms or changes in the number of infected plants among the *ap2m-1* mutant, the complementation line and wt at 14 dpi, indicating that the absence of AP2M does not impair systemic infection of OuMV (Table S3).

### 4. Y and LL motif MP mutants display changes in subcellular localization over time

We next examined whether Y and LL motif MP mutants display different intracellular localization patterns over time (Fig. S5). To characterize the subcellular distribution of MP mutants, we transiently expressed GFP:MPwt (Fig. S5A), as a control, and GFP:MP mutants (Fig. S5B-F) with RdRp and CP in *N. benthamiana* epidermal cells. GFP-tagged Y motif MP mutants, GFP:MP_Y and GFP:MP_Y/G, appeared to be distributed in the cytoplasm at 36 hpi and 48 hpi (Fig. S5B-C, left panels) without any noticeable change at 72 hpi (Fig. S5B-C, right panel). Y motif mutants accumulated in transvacuolar strands and in patches at the cell cortex. Puncta that were consistently observed in GFP:MPwt (Movie S2) were no longer apparent in GFP:MP_Y and GFP:MP_Y/G (Movie S3). Occasionally, a few punctate structures were observed in GFP:MP_Y/G; however, these structures were either stationary or had rather restricted movement (Movie S3). GFP-tagged LL motif MP mutants had a cytoplasmic localization at earlier time points (Fig. S5D-F), with peripheral and cytoplasmic puncta readily observable at 60 hpi, similar to GFP:MPwt (Fig. S5D-F, Movie S4).

### 5. Mutations in the Y but not in the LL motif impair plasma membrane targeting of MP

To investigate whether Y and LL motif MP mutants are targeted to the plasma membrane, we stained *N. benthamiana* protoplasts expressing GFP:MPwt or GFP:MP mutants with FM4-64, a plasma membrane marker. GFP:MPwt showed a continuous signal at the cell periphery and this signal strongly overlapped with FM4-64 (Fig. 3A). The peripheral signal of GFP:MP_Y and GFP:MP_Y/G often did not co-localize with FM4-64 (Fig. 3B-C). A similar pattern was observed in *N. benthamiana* epidermal cells expressing GFP:MP_Y or GFP:MP_Y/G, where the peak of FM4-64 often positioned in between two peaks of the MP mutants, suggesting that plasma membrane localization of GFP:MP_Y and GFP:MP_Y/G followed a different pattern than GFP:MPwt (Fig. S6). Moreover, both mutants exhibited an intense and diffuse cytoplasmic signal. The peripheral signal of GFP:MP_D/G mostly overlapped with FM4-64 (Fig. 3D). The movement deficient mutant GFP:MP_D also displayed a peripheral signal that co-localized with FM4-64; however, the GFP signal was not evenly distributed along the plasma membrane and accumulated at specific sites (Fig. 3E). GFP:MP_LI/GG, on the other hand, had regions where the signal is either continuous or puncta (Fig. 3F).

**Figure 3.**
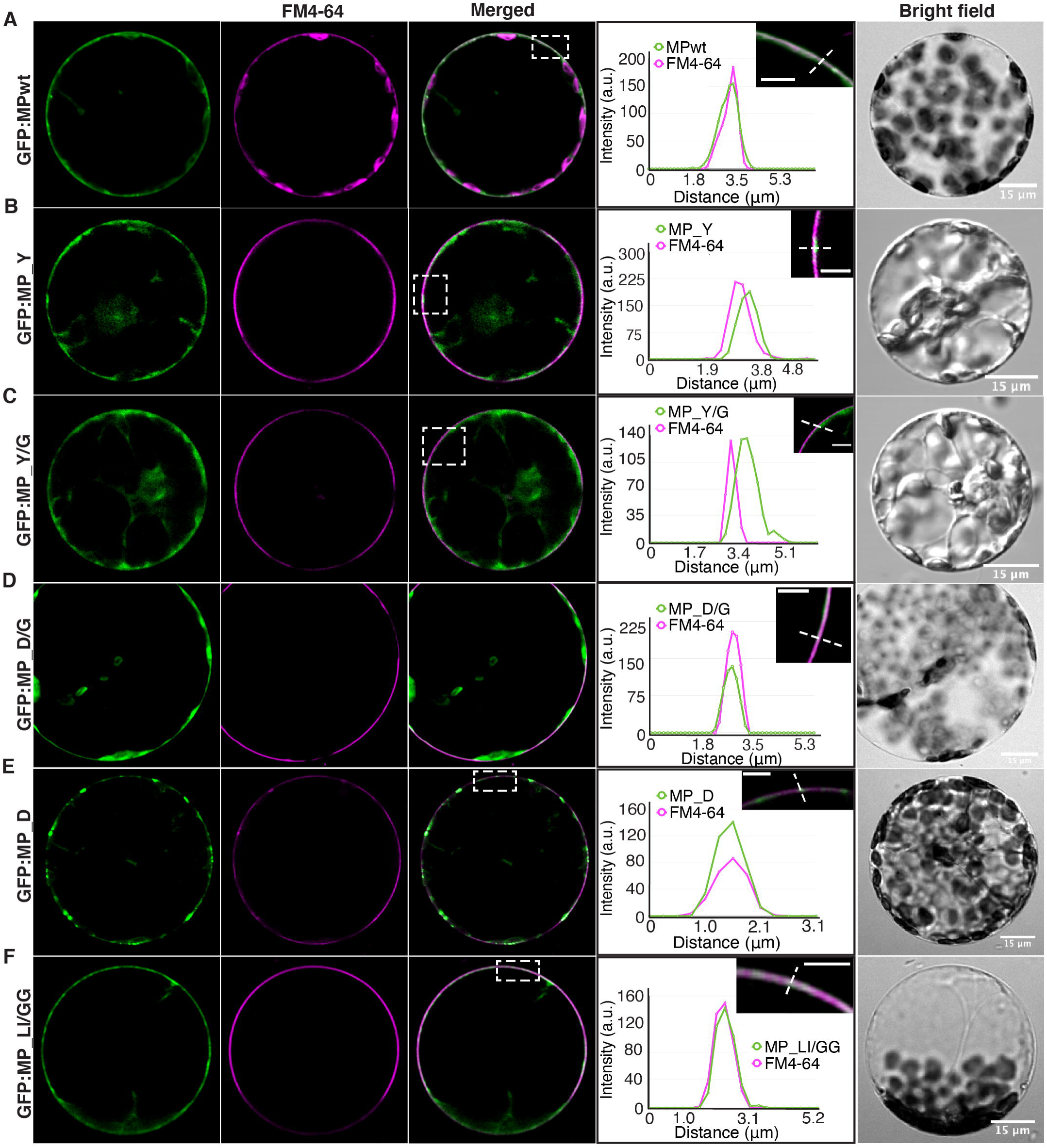
Plasma membrane localization of MP mutants. *N. benthamiana* protoplasts expressing **(A)** GFP:MPwt or GFP:MP mutants, **(B)** GFP:MP_Y, **(C)** GFP:MP_Y/G, **(D)** GFP:MP_D/G, **(E)** GFP:MP_D and **(F)** GFP:MP_LI/GG along with RdRp and CP. Protoplasts were stained with FM4-64, a plasma membrane marker (magenta). Selected regions (white rectangles) showing the intensity profile of the diagonal line were generated by ImageJ. Images represent single optical planes. Scale bar, 5 μm (magnified region).

To investigate these observations, *N. benthamiana* epidermal cells expressing GFP:MPwt or mutants were subjected to plasmolysis. While GFP:MPwt, GFP:MP_D, GFP:MP_D/G and GFP:MP_LI/GG were associated with Hechtian strands that attach the plasma membrane to the cell wall, GFP:MP_Y or GFP:MP_Y/G did not exhibit the same pattern, as GFP-marked strands were absent from the space between the protoplast and the cell wall (Fig. 4). Plasma membrane localization of GFP:MP_D and GFP:MP_LI/GG suggests that mutations of the LL motif do not impair plasma membrane targeting of MP but alter uniform distribution of MP along the plasma membrane.

**Figure 4.**
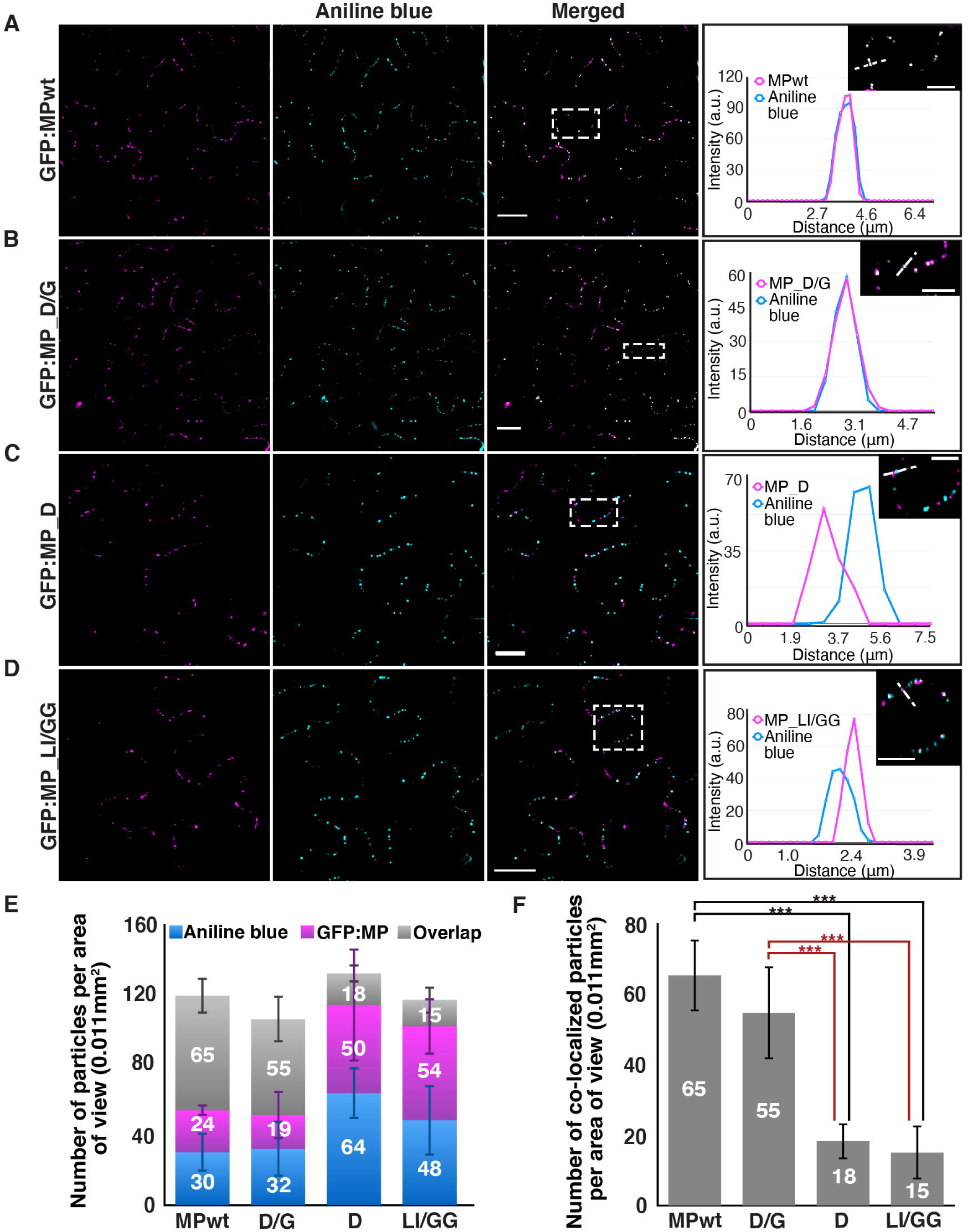
LL motif MP mutants are targeted to the plasma membrane. *N. benthamiana* epidermal cells expressing GFP:MPwt (top left panel) or GFP:MP mutants, GFP:MP_Y (top middle panel), GFP:MP_Y/G (top right panel), GFP:MP_D/G (bottom left panel), GFP:MP_D (bottom middle panel), and GFP:MP_LI/GG (bottom right panel) along with RdRp and CP. were plasmolyzed with 1M mannitol. Arrows show the retracted plasma membrane and asterisks denote Hechtian strands. Images were taken at 62-66 hpi and represent single optical planes. Scale bar, 10 μm.

### 6. Movement deficient LL motif MP mutants failed to localize to PD

The punctate localization of the two movement-deficient LL motif MP mutants, MP_D and MP_LI/GG, along the cell periphery raises the question of whether these punctate structures are associated with PD. We examined the targeting of MP mutants to PD by staining *N. benthamiana* epidermal cells expressing GFP:MPwt or MP mutants with aniline blue, a callose-specific dye. Consistent with our previous results (37), GFP:MPwt co-localized with callose deposits at PD (Fig. 5A). GFP:MP_D/G, which is able to support movement, exhibited a similar localization pattern compared to GFP:MPwt, and the intensity profile across the PD strongly coaligned with that of aniline blue (Fig. 5B). Although peripheral punctate structures marked by GFP:MP_D and GFP:MP_LI/GG did not predominantly localize to callose deposits (Fig. 5C), GFP:MP_LI/GG labeled structures were in close proximity to PD (Fig. 5D). The quantification of points labeled by GFP:MPwt or GFP:MP mutants and aniline blue revealed that some of GFP:MP_D and GFP:MP_LI/GG co-localize with callose deposits, but the number of co-localized signal was much lower than that of GFP:MPwt and GFP:MP_D/G (Fig. 5E). The statistical analysis of co-localization revealed significant differences between GFP:MPwt or GFP:MP_D/G and movement deficient MP mutants, GFP:MP_D and GFP:MP_LI/GG (Fig. 5F). In contrast, no significant difference between GFP:MPwt and GFP:MP_D/G was observed (Fig. 5F). These results suggest that OuMV movement correlates with the ability of its movement protein to be targeted to PD. Results for all mutants are summarized in Table S4.

**Figure 5.**
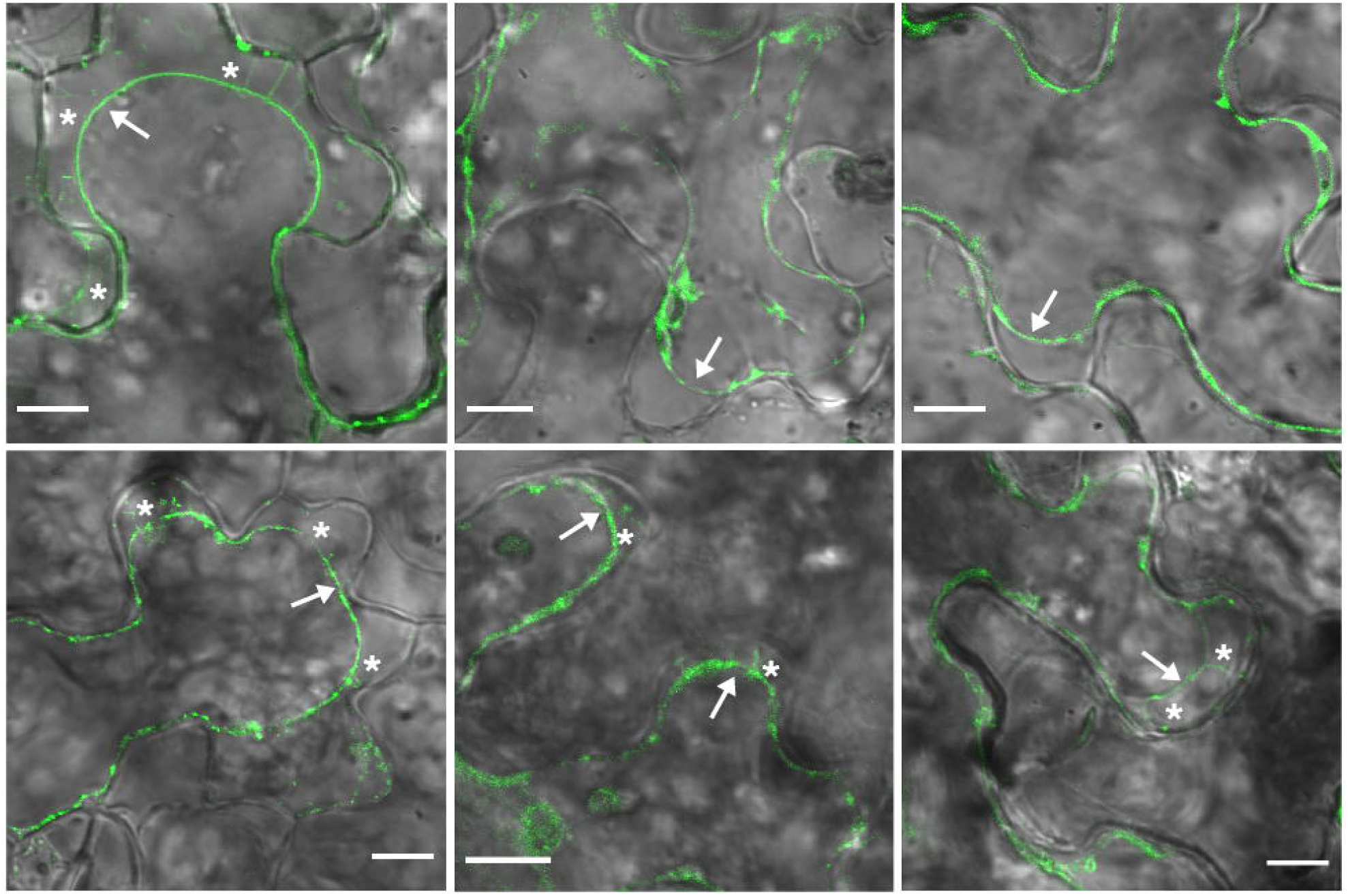
Movement deficient LL motif MP mutants do not localize to callose deposits. N. *benthamiana* epidermal cells expressing **(A)** GFP:MPwt or GFP:MP mutants, **(B)** GFP:MP_D/G, **(C)** GFP:MP_D, and **(D)** GFP:MP_LI/GG (magenta) with RdRp and CP. Aniline blue was used to mark callose deposits (cyan). Selected regions (white rectangles) show the intensity profile of the diagonal line generated from images that were adjusted to a lower intensity by ImageJ. Images represent single optical planes. Scale bar, 20μm, 10 μm (magnified region). **(E)** The quantification of callose deposits (blue), punctate structures labeled by OuMV GFP:MPwt or GFP:MP mutants (magenta), and co-localized particles (gray) per area of view. The particles were counted using the ComDet v.0.4.1 plugin in ImageJ. At least 6 areas of view were examined (n≥6) and the total number of counted particles for each sample is more than 700. Stacked bar graph represents mean±standard deviation **(F)** The statistical analysis of co-localized particles in (E). Bar graph represents PD-localized OuMV GFP:MPwt or GFP:MP mutants per area of view (mean± standard deviation, n≥6). ***, P < 0.001 by ANOVA, followed by Tukey’s HSD.

To understand the specific role of LL motif in the plasmodesmata targeting of OuMV MP, N-terminal domain truncated OuMV MP mutants retaining the LL motif, MP_84-first 84 amino acids of MP, and retaining both LL and Y motifs, MP_102-first 102 amino acids of MP, were generated based on predicted secondary structure of MP. We monitored the targeting of truncated MP as described above. Both mutants were, however, unable to reach PD (Fig. 6), indicating that the presence of the LL motif in a truncated MP variant is not sufficient to support plasmodesmata targeting of MP, and the C-terminal region of MP is also involved in this process. Moreover, MP_102 had noticeably lower expression levels, close to the background signal, compared to MP_84.

**Figure 6.**
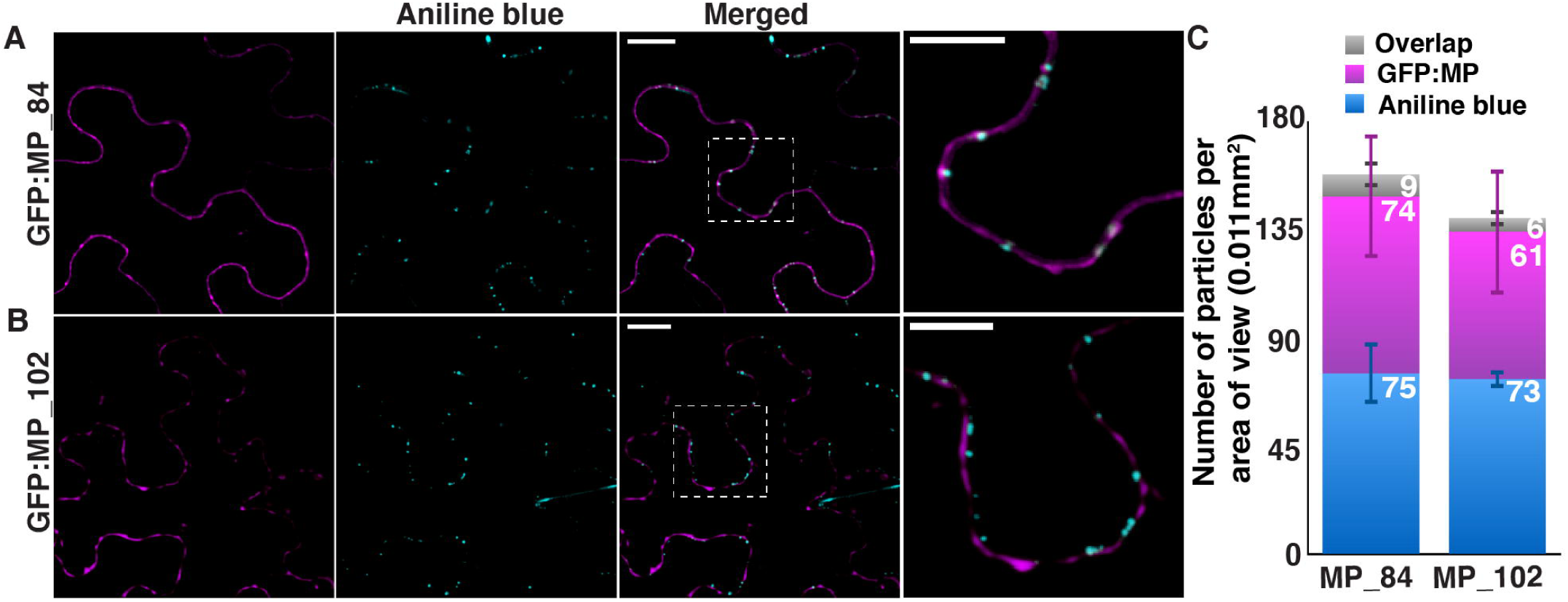
Truncated OuMV MP mutants retaining sorting motifs failed to localize to callose deposits. *N. benthamiana* epidermal cells expressing **(A)** GFP:MP_84, **(B)** GFP:MP_102 (magenta) with RdRP and CP. Aniline blue was used to mark callose deposits (cyan). Images represent single optical planes. Scale bar, 15μm, 10 μm (magnified region). **(C)** The quantification of callose deposits (blue), particles labeled by OuMV GFP:MP truncated mutants and co-localized particles (gray) per area of view. The particles were counted using the ComDet plugin in ImageJ. Four areas of view were examined (n=4) and the total number of counted particles for each sample is more than 600. Stacked bar graph represents mean±standard deviation.

## Discussion

In this study, we investigated the intracellular trafficking of OuMV MP in the context of post-Golgi transport routes and the function of Y and LL sorting motifs. Despite the complexity of plant protein sorting arising from the fact that TGN acts as the main sorting hub for Golgi-derived cargoes and internalized cargoes from the plasma membrane, we were able to partially unravel the trafficking routes that OuMV MP hijacks by using organelle markers in combination with inhibitors of vesicle trafficking. TGN localization of OuMV MP, together with its co-localization with FM4-64 stained compartments, demonstrates that OuMV MP trafficks in endosomal pathways (Fig. 1, A-B). Co-localization of endosomal compartments and MPs of other viruses has been reported before (16,34); however, this is the first study showing that a viral MP resides in the TGN, extending knowledge of the involvement of post-Golgi trafficking pathways in virus movement. We also showed that the canonical secretory pathway is not involved in the transport of OuMV MP since the MP did not co-localize with the Golgi in the absence or presence of BFA (Fig. 1D). Moreover, the redistribution of OuMV MP into the same smaller vesicles as the TGN marker upon low dose BFA treatment suggests that MP exhibits BFA sensitivity only at the TGN, but not at the Golgi. Dependence on the canonical secretory pathway varies between plant viruses (4). For example, whereas the secretory pathway is not involved in targeting of the CPMV MP to the cell periphery (51), a functional secretory pathway is necessary for the correct localization of the GFLV MP (14). In addition, OuMV MP was not associated with MVBs/PVCs in the absence or presence of Wm (Fig. 1C), confirming that MP trafficks in the first stages of vesicular trafficking, but not in MVB/PVC-to-vacuole pathway, in contrast to CaMV MP (16) and PTMV TGB2 (34), which co-localize with ARA7-positive MVBs/PVCs. Although OuMV MP showed sensitivity to Wm, this response was different from that of MVBs/PVCs, probably due to the effect of Wm on endocytosis. Based on the finding that OuMV MP localizes to the TGN and does not enter either the canonical secretory or MVB/PVC-to-vacuole pathway, we hypothesize that MP constitutively cycles between the plasma membrane and TGN. Although a similar recycling mechanism has been proposed for CaMV MP (16), the TGN localization of CaMV MP was not explored.

Glycine substitutions in the Y motif altered the intracellular distribution of MP. A key question emerging from this observation is whether the Y motif in OuMV MP is a functional internalization motif. Y motifs are recognized by μ-subunits of APs 1-4 (24,52,53). In plants, clathrin-mediated endocytosis (CME) of plasma membrane proteins is achieved by the recognition of Y motifs in their cytoplasmic domains by AP2M (54–57), and it has been previously shown that CaMV MP is able to bind Arabidopsis AP2M through three Y motifs (16). While mutations in these motifs do not affect targeting of the protein to the plasma membrane, its internalization relies on at least one functional Y motif. Unlike CaMV, mutations in the Y motif retained OuMV MP in the cytoplasm. Additionally, we did not observe any changes in the infectivity of OuMV in the absence of AP2M in Arabidopsis (Table S3). It is, however, known that AP2M is not the sole regulator of CME in plants; a CME adaptor complex, TPLATE complex (TPC), is involved in the initiation of CME and is thus crucial for recruitment of clathrin and AP2 (58). Moreover, AP2 subunits are considered to form partially active complexes in the absence of single subunits (59). Hence, other AP2 subunits and/or TPC may be important for internalization of OuMV MP, and in turn for infectivity of OuMV. Indeed, the identification of AP2α as a candidate interactor (Table S2) suggests that the internalization of OuMV MP may still occur via an AP2-dependent mechanism. Alternatively, Y motif may have other functions such as maintaining correct protein conformation. Glycine substitutions into the Y motif of OuMV MP may induce conformational changes, leading to masking of the plasma membrane targeting signal. In PSLV and PMTV, the Y motif is required for the delivery of viral proteins to the cell periphery and PD, but this motif is important for conformation of the protein, rather than its internalization (35). We previously identified three amino acid residues (98, 150 and 169) within the core region of OuMV MP (63-206), which are responsible for localization of MP at the cell periphery, and these residues are possibly involved in the correct folding/function of OuMV MP (37). The Y motif of OuMV MP is located in the core region of the MP, suggesting that it may have a structural role, similar to the one seen for PSLV TGB3 and PMTV TGB3. Moreover, The Y motif of OuMV MP may be important for recruiting viral and cellular factors that are necessary for intracellular targeting as the YQDLN motif in the TGB3 of PSLV plays a crucial role in its intracellular transport, facilitating the formation of higher-order complexes (60).

OuMV MP has the classical [D/E]XXXL[L/I] sorting motif in which an aspartic acid is located at the position 1. Replacing the aspartic acid with glycine in DPAILI does not affect the localization of the MP and the resulting mutant behaves similar to the wild-type virus (Fig. S5E, Fig. 3-4D; Fig. 5B). Conversely, substitutions of leucine and isoleucine with a pair of glycine residues impair the transport of the OuMV MP to PD. In plants, the LL motif is recognized by AP complexes (61,62). The LL motif of the tonoplast-localized ion transporter VTI1, EKQTLL, exhibits affinity for AP1γ1/2 and σ1/2 subunits (61). Notably, AP1γ co-purified with OuMV MP (Table S2), thus strengthening the possibility of AP1γ as an interacting partner of OuMV MP. Intriguingly, AP2β has been recently shown to interact with turnip mosaic virus (TuMV) proteins (i.e., 6K2, VPg, CI, and CP), which harbor putative LL motifs, implicating a potential role of the LL motif in the endocytosis during viral infection (63).

Previous studies have shown that viral proteins traffic in endocytic and vacuolar pathways (11,16,34,63,64). Our results support the hypothesis that plant viral proteins take advantage of post-Golgi trafficking pathways to facilitate cell-to-cell movement and provide new insights into these pathways. Based on our findings, we propose the following model (Fig. 7): OuMV MP is targeted to the plasma membrane through a mechanism that is independent of the conventional secretory pathway. While MPs of some viruses reach the plasma membrane and PD through the ER (4,65), we do not have evidence at this point that OuMV MP follows this model. Once OuMV MP reaches the plasma membrane, it is endocytosed, sorted into the TGN, and then recycled back to the plasma membrane, bypassing the MVB/PVC-to-vacuole pathway, to reach PD. The retrieval of MP from the plasma membrane does not appear to depend on the Y motif.

**Figure 7.**
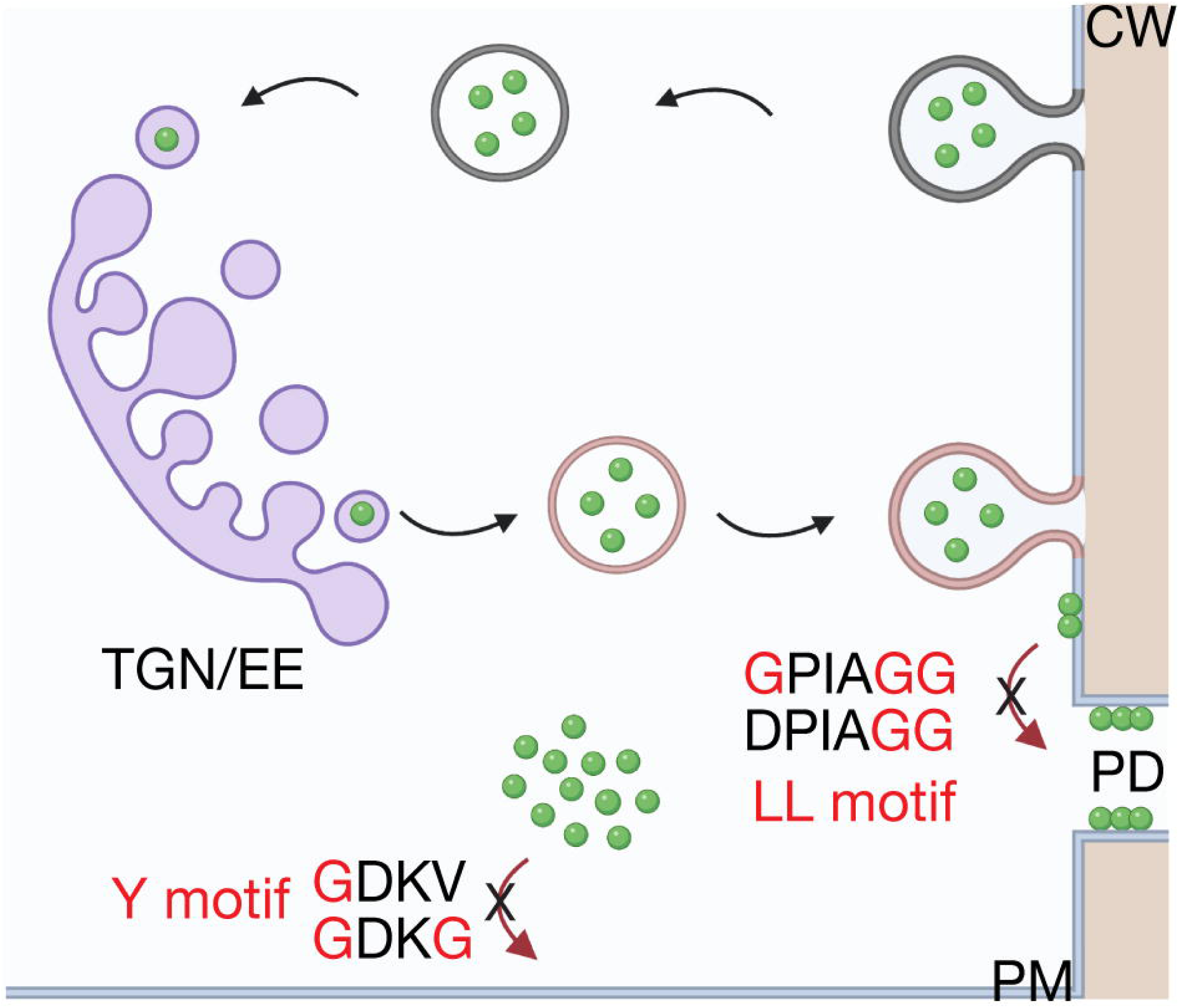
Working model for intracellular trafficking of OuMV MP. The MP (green) is targeted to the plasma membrane via an unknown mechanism. It is retrieved from the plasma membrane (dark gray) and transported to the TGN (purple). From the TGN, the MP is recycled back to the plasma membrane (pink). The Y motif possibly acts as a structural motif rather than a functional internalization motif, whereas the LL motif is required for correct plasmodesmata targeting of MP. TGN/EE, trans-Golgi network/early endosome; PM, plasma membrane; PD, plasmodesmata; CW, cell wall. The model was created with BioRender.com

On the other hand, the PD targeting of MP relies on the LL motif, potentially through its function in endocytic pathways. The precise molecular mechanism of MP targeting is, however, still unknown. In fact, the N-terminal region of MP retaining the LL motif does not act as a PD-targeting signal (Fig. 6), as opposed to the TMV MP, in which the first 50 amino acid residues were characterized as a PD localization signal (66). Moreover, the C-terminus of OuMV has a predicted disordered region with many potential phosphorylation sites that may be important for its regulation and interaction with host proteins. The model presented in this study requires further examination to understand the mechanistic details of the targeting of OuMV MP to PD. Movement proteins are often associated with the sites of viral replication. Thus, it is extremely hard to untangle if a disturbance in viral replication, as seen with our mutants, is due to mislocalization of the viral MP from the replication site or to modifications of the sites of replication. Nevertheless, our characterization of OuMV MP-targeted pathways partially reveals missing molecular and cellular components of OuMV movement and virus-host interactions.

## Materials and Methods

### Plant material and growth conditions

*N. benthamiana* plants were grown in a growth chamber set at 25°C with a 16 h photoperiod and used at 4-5 weeks old. *Arabidopsis thaliana* seeds were surface sterilized using 50% bleach solution for 5 minutes, rinsed five times with sterile distilled water and plated on ½ MS plates containing 1% agar. Ten-day old seedlings were transferred to soil, grown in a growth chamber at 22 °C, 60% relative humidity, and at a light intensity of 120 µmol/m^2^s^1^ at a 10 h photoperiod, and used when 4-5 weeks old.

*Arabidopsis thaliana* ecotype Col-0 was used for infectivity assays. The AP2 knockout mutant *ap2m-1* (SALK_083693) and AP2M-YFP complementation line (67) were provided by Dr. Ying Gu (The Pennsylvania State University, University Park, PA, USA) and genotyped for homozygosity using the primers in Table S5.

### Plasmid constructions and *Agrobacterium-*mediated transient expression in plant leaves

The OuMV infectious clone, consisting of the three plasmids pGC-RNA1 (RdRp), pGC-RNA2 (MP) and pGC-RNA3 (CP), and GFP:MP plasmid has been previously described (37,38). Mutations were inserted into MP and GFP:MP plasmid with the QuickChange Site-Directed Mutagenesis Kit (Stratagene, La Jolla, CA, USA) using primers in Table S5.

The Golgi marker, Man49:mCherry (CD3-967) (48), the TGN marker, mCherry:VTI12 (CD3-781541) (42), and the MVB/PVC marker, mCherry:RHA1 (CD3-781597) (42) were obtained from ABRC (http://www.arabidopsis.org/). The fragments of Ubq10pro::mCherry::VTI12 and Ubq10pro::mCherry::RHA1 were amplified from the original plasmids using the primers in Table S5, and inserted into XhoI/SpeI-digested pORE O2 (CD3-921) (68). All plasmids were transformed into *A. tumefaciens* strain C58C1.

*A. tumefaciens* cultures carrying the desired plasmids were grown in a shaking incubator at 28°C in 2xYT medium supplemented with 50 µg/ml kanamycin and 5 µg/mL tetracycline. Cultures were pelleted at an optical density at 600 nm (OD600) of 1.0 and resuspended in agroinfiltration buffer (10 mM MES, 10 mM MgCl_2_ and 100 µM acetosyringone). Bacterial suspensions were generated by mixing the desired combinations at a final optical density at 600 nm (OD600) of 0.2 and 0.05 for GFP:MP/RdRp/CP and the organelle markers, respectively. The mixtures were incubated at room temperature for 2 h in the dark and infiltrated on the abaxial side of fully expanded leaves using a needleless syringe.

### Infectivity assays, ELISA, and RT-PCR

Viral infections were started either from agroinfiltration of a mixture of *Agrobacterium* cultures carrying the three plasmids mentioned above or by mechanical inoculation using sap obtained by grinding virus-infected fresh *N. benthamiana* tissue in 0.05 M phosphate buffer supplemented with carborundum and silica powders (37). Six *N. benthamiana* and 12 Arabidopsis plants were used for infectivity assays. Double antibody sandwich-enzyme linked immunosorbent assay (DAS-ELISA) was performed as previously described (37). Leaves were sampled for ELISA testing at 14 dpi. Experiments were repeated three times.

RNA extraction and RT-PCR were performed as described in (37). Total RNA (100 ng) was used for RT-PCR, and full-length coding sequence of MP was amplified using primers listed in Table S5.

### Replication assay

The effect of specific amino acid substitutions in MP on virus replication was measured by agroinfiltration of local leaves with pGC-RNA2 (MP) or mutants, together with pGC-RNA1 (RdRp), pGC-RNA3 (CP), and pBin61-GFP at an OD600 of 1.0. The GFP signal at the infiltrated sites was monitored at 36 hpi to confirm that agroinfiltrated areas were saturated. Two leaf spots from 3 leaves were pooled for each sample at 24 hpi and 48 hpi. The virus accumulation was measured by quantifying RNA1 by qPCR using primers listed in Table S5. The qPCR was performed in duplicates using iTaq™ Universal SYBR® Green Supermix (BioRad) and a CFX96 (BioRad), following the manufacturer’s instructions. The 2^-ΔΔCT^ method was used to assess fold changes in replication of wild-type and mutants. The data was normalized by first using 18S ribosomal RNA and then a wild-type control at 24 hpi showing the lowest expression. Four plants were used for qPCR analysis. Statistical analysis was performed using Student’s t-test.

### Protoplast isolation from infected leaves

Protoplasts were prepared from leaves of *N. benthamiana* expressing GFP:MPwt or GFP:MP mutants (together with RdRp and CP) at 66 hpi. Leaf sections were cut into small fragments (∼1 cm^2^) using a razor blade and incubated in the enzyme solution on a rotor shaker for 6 h at room temperature, as previously described (37).

### Plasmolysis, inhibitor, and staining experiments

Plasmolysis experiments were carried out by treating *N. benthamiana* leaf sections expressing GFP:MP or mutants, together with RdRp and CP, with 1 M mannitol for 45 min. The leaf sections were imaged at 62 – 66 hpi.

*N. benthamiana* leaves were infiltrated with 10 μM BFA (stock: 10mg/ml in DMSO; MilliporeSigma) or 33 μM Wortmannin (stock: 10 mM in DMSO; MilliporeSigma). Equivalent amounts of DMSO were used as controls. Leaf samples were imaged after 1 h of the indicated treatment, as described below. *N. benthamiana* protoplasts and epidermal cells were stained with the dye FM4-64 (Thermo Fisher Scientific) at a final concentration of 10 μM (stock: 1 mM in water) for 5-15 min. For staining callose deposits, *N. benthamiana* leaves were infiltrated with aniline blue (MilliporeSigma) at a final concentration of 0.1% (stock: 1% (v/w) in 50 mM potassium phosphate buffer, pH 9.0) and incubated for 30 min. All experiments were repeated at least twice.

### Confocal microscopy and quantification

Leaf sections or protoplasts were mounted on a microscope slide (Thermo Fisher Scientific) covered with a #1.5 coverslip (Corning). For confocal microscopy, images were acquired with a Zeiss Cell Observer SD microscope equipped with a Yokogawa CSU-X1 spinning disk head and Plan-Apochromat 63X/NA 1.4 and 40X/NA 1.3, and an Olympus Fluoview (FV) 1000 microscope with a Plan-Apochromat 40X/NA 1.15 water immersion and 60X/NA 1.4 oil immersion objectives. A 543-nm excitation laser line and an emission bandpass filter 585 – 660 nm (Olympus FV1000) were used for imaging of FM-4-64. Aniline blue was imaged using 405-nm excitation laser with an emission bandpass filter 425 – 475 nm (Olympus FV1000 and Zeiss Cell Observer SD). Z-stacks of optical sections were collected at 0.2 μm intervals for time-dependent subcellular localization experiments and at 0.5 μm intervals for the rest of the experiments. Images were analyzed and processed using ImageJ (http://imagej.nih.gov/ij. The background was subtracted, and the signal intensity was adjusted. Corresponding images were later processed by Adobe Photoshop CC 2017. All images were assembled in Adobe Illustrator CC 2022.

Callose deposits were quantified using the ComDet v.0.4.1 plugin in ImageJ. The particle detection was performed by setting particle size to 2 pixels. For co-localization experiments, maximum distance between spots was adjusted to 2 pixels. Statistical analysis was performed using ANOVA.

### Protein purification

The coding sequence of OuMV MP was cloned into in pMAL-c4x (New England Biolabs; kindly provided by Dr. Gitta Coaker (University of California-Davis, CA, USA), in frame with MBP. MBP-MP was expressed in BL21 (DE3), and the expression was induced with 0.3 mM IPTG for 2 h at 37° C. The pellets were resuspended with amylose column buffer (20 mM Tris-HCl pH7.5, 0.2 M NaCl, 1 mM EDTA) supplemented with 1X Halt protease inhibitor cocktail (Thermo Fisher Scientific) and sonicated five times for 10 sec. The soluble fraction of MBP-MP was applied to pre-equilibrated amylose agarose resin (New England Biolabs). After washing the column three times with amylose column buffer, MBP-MP was eluted with five fractions of amylose column buffer supplemented with 30 mM maltose. The fraction containing the highest amount of MBP-MP was concentrated using an Amicon Ultra-15 Centrifugal Filter Unit with Ultracel-50kDa (MilliporeSigma) and stored in storage buffer (20 mM HEPES pH 7.5, 5 mM MgCl_2_, 1 mM EDTA, 1 mM DTT, 20% glycerol).

### Co-IP Assay

One gram of five-weeks old Arabidopsis rosette leaves were homogenized in 2ml of cold IP buffer (20 mM HEPES pH 7.4, 100 mM NaCl, 5 mM MgCl_2_, 5% glycerol, 0.5% Triton X-100, and 1X Halt protease/phosphatase inhibitor cocktail, Thermo Scientific) using a pre-cooled mortar and pestle. The homogenate was first centrifuged at 1,000g for 10 min at 4°C, and then at 10,000g for 10 min at 4°C. For the second part of the study, 10 grams of five-weeks old Arabidopsis rosette leaves were homogenized in 15 ml of cold homogenization buffer (50 mM HEPES pH 7.4, 330 mM sucrose, 10% glycerol and 1X Halt protease/phosphatase inhibitor cocktail, Thermo Scientific) using a pre-cooled mortar and pestle and filtered through four layers of Miracloth. The homogenate was first centrifuged at 1,000g for 10 min at 4°C, and then at 10,000g for 10min at 4°C. The supernatant was centrifuged to pellet microsomal membranes at 100,000g for 1 h at 4°C. The supernatant was used for the assay. Five μg of MBP or MBP-MP were bound to MBP-Trap-A beads (ChromoTek) for 1 h at 4°C with rotation and washed three times with washing buffer (10 mM Tris-HCl pH 7.5, 150 mM NaCl, 5 mM MgCl_2_, 0.5 mM EDTA) and two times with IP buffer. MBP or MBP-MP bound beads were incubated with plant lysate for 2 h at 4°C with rotation. Samples were washed three times with an IP buffer with 0.2% Triton X-100 and twice with 50 mM (NH_4_)_2_CO_3_, pH 8.0 to remove the detergent.

### On-bead trypsin digestion and LC-MS/MS

The spin columns were blocked with plugs and 100 μl of on-bead trypsin digestion buffer (50 mM (NH4)2CO3, 0.01% ProteaseMax (Promega), and 1 μg Trypsin (Promega) was added to the columns. The samples were digested for 1 h at 37°C with gentle shaking to allow beads to be mixed in the solution. The digested samples were collected in low-bind tubes (Eppendorf). The beads were washed twice with 50 μl of on-bead trypsin digestion wash buffer (50 mM (NH_4_)_2_CO_3_, 0.01% ProteaseMax, and 5 mM DTT). The samples were incubated for 30 min at 37°C. Five μl of 500 mM Iodoacetamide (IAA) were added to digested samples and the samples were incubated at 37°C overnight in the dark. Three μl of 1M DTT were added to the samples to quench IAA and incubated for 30 min at 37°C. Ten ul of 10% trifluoroacetic acid (TFA) were added to a final concentration of 0.5% to acidify the samples and stop the digestion. The samples were incubated for 5 min at room temperature. Samples were snap-frozen in liquid nitrogen and lyophilized. Lyophilized samples were sent to the Penn State College of Medicine’s Mass Spectrometry & Proteomics Core for protein identification. The samples were desalted with C18 stage tips and injected into an ABSciex TripleTOF 5600+ (Applied Biosystems). MS/MS data were analyzed using ProteinPilot software. Unused Score > 1.3 (95% confidence) was used as a cut-off threshold to identify proteins.

### Protein Lipid Overlay Assay

The phospholipid-binding analysis of the MBP-MP was performed using PIP Strips (P-6001; Echelon Biosciences). The PIP strips were blocked for 1 h at room temperature in Tris-buffered saline (TBS) with 4% non-fat milk and then incubated 1 h at room temperature in blocking buffer containing 3.5 µg/ml MBP-MP. The membranes were washed with TBS and incubated with a horseradish peroxidase (HRP)-conjugated anti-MBP monoclonal antibody (New England Biolabs; 1:5000).

### Protein extraction, SDS–PAGE, immunoblotting

Leaf sections were collected from agroinfiltrated leaves and upper uninoculated leaves and homogenized in a 2x SDS-PAGE buffer. Extracted proteins were separated in SDS polyacrylamide gels (4% stacking gel, 10% separation gel). The proteins were transferred to PVDF membranes (MilliporeSigma) for western blot analysis. OuMV MP was detected using a non-commercial anti-MP (A314) antiserum as described in (36), followed by incubation with an anti-rabbit HRP-conjugated secondary antibody (NXA931; MilliporeSigma) diluted at 1:20000 (v/v). SuperSignal™ West Femto Maximum Sensitivity Substrate (Thermo Fisher Scientific) was used for detecting HRP activity.

## Acknowledgements

This work was supported by USDA HATCH Accession No: 1009992, Project No. PEN04604 to C. Rosa and a grant of Penn State College of Agricultural Sciences to N. Ozber.

## Notes

### Competing Interest Statement

The authors have declared no competing interest.

## References

1. Benitez-Alfonso Y, Faulkner C, Ritzenthaler C, Maule AJ. Plasmodesmata: Gateways to Local and Systemic Virus Infection. Mol Plant-Microbe Interact (2010) 23:1403–1412. doi: 10.1094/MPMI-05-10-0116

2. Niehl A, Heinlein M. Cellular pathways for viral transport through plasmodesmata. Protoplasma (2011) 248:75–99. doi: 10.1007/s00709-010-0246-1

3. Wu X, Cheng X. Intercellular movement of plant RNA viruses: Targeting replication complexes to the plasmodesma for both accuracy and efficiency. Traffic (2020) 21:725–736. doi: https://doi.org/10.1111/tra.12768

4. Pitzalis N, Heinlein M. The roles of membranes and associated cytoskeleton in plant virus replication and cell-to-cell movement. J Exp Bot (2018) 69:117–132. doi: 10.1093/jxb/erx334

5. Kumar G, Dasgupta I. Variability, Functions and Interactions of Plant Virus Movement Proteins: What Do We Know So Far? Microorg (2021) 9: doi: 10.3390/microorganisms9040695

6. McLean BG, Zupan J, Zambryski PC. Tobacco mosaic virus movement protein associates with the cytoskeleton in tobacco cells. Plant Cell (1995) 7:2101–2114. doi: 10.1105/tpc.7.12.2101

7. Jamie A, Emmanuel B, Mark S, Adrian S, Christophe R, Manfred H. Tobacco Mosaic Virus Movement Protein Functions as a Structural Microtubule-Associated Protein. J Virol (2006) 80:8329–8344. doi: 10.1128/JVI.00540-06

8. Kragler F, Curin M, Trutnyeva K, Gansch A, Waigmann E. MPB2C, a Microtubule-Associated Plant Protein Binds to and Interferes with Cell-to-Cell Transport of Tobacco Mosaic Virus Movement Protein. Plant Physiol (2003) 132:1870–1883. doi: 10.1104/pp.103.022269

9. Brandner K, Sambade A, Boutant E, Didier P, Mély Y, Ritzenthaler C, Heinlein M. Tobacco Mosaic Virus Movement Protein Interacts with Green Fluorescent Protein-Tagged Microtubule End-Binding Protein 1. Plant Physiol (2008) 147:611–623. doi: 10.1104/pp.108.117481

10. Chen M-H, Tian G-W, Gafni Y, Citovsky V. Effects of Calreticulin on Viral Cell-to-Cell Movement. Plant Physiol (2005) 138:1866–1876. doi: 10.1104/pp.105.064386

11. D. Lj, G. Ls. Arabidopsis synaptotagmin SYTA regulates endocytosis and virus movement protein cell-to-cell transport. Proc Natl Acad Sci (2010) 107:2491–2496. doi: 10.1073/pnas.0909080107

12. Tran P-T, Citovsky V. Receptor-like kinase BAM1 facilitates early movement of the Tobacco mosaic virus. Commun Biol (2021) 4:511. doi: 10.1038/s42003-021-02041-0

13. Ana P, Luis M-G, Silvia T, Vicente P A. S-NJ, Ismael M A. S. The Tobacco mosaic virus Movement Protein Associates with but Does Not Integrate into Biological Membranes. J Virol (2014) 88:3016–3026. doi: 10.1128/JVI.03648-13

14. Laporte C, Vetter G, Loudes A-M, Robinson DG, Hillmer S, Stussi-Garaud C, Ritzenthaler C. Involvement of the Secretory Pathway and the Cytoskeleton in Intracellular Targeting and Tubule Assembly of Grapevine fanleaf virus Movement Protein in Tobacco BY-2 Cells. Plant Cell (2003) 15:2058–2075. doi: 10.1105/tpc.013896

15. Huang Z, Andrianov VM, Han Y, Howell SH. Identification of Arabidopsis proteins that interact with the cauliflower mosaic virus (CaMV) movement protein. Plant Mol Biol (2001) 47:663–675. doi: 10.1023/A:1012491913431

16. Carluccio AV, Zicca S, Stavolone L. Hitching a Ride on Vesicles: Cauliflower Mosaic Virus Movement Protein Trafficking in the Endomembrane System. Plant Physiol (2014) 164:1261–1270. doi: 10.1104/pp.113.234534

17. Amari K, Boutant E, Hofmann C, Schmitt-Keichinger C, Fernandez-Calvino L, Didier P, Lerich A, Mutterer J, Thomas CL, Heinlein M, et al. A family of plasmodesmal proteins with receptor-like properties for plant viral movement proteins. PLoS Pathog (2010) 6:e1001119–e1001119. doi: 10.1371/journal.ppat.1001119

18. Paape M, Solovyev AG, Erokhina TN, Minina EA, Schepetilnikov M V, Lesemann D-E, Schiemann J, Morozov SY, Kellmann J-W. At-4/1, an Interactor of the Tomato spotted wilt virus Movement Protein, Belongs to a New Family of Plant Proteins Capable of Directed Intra-and Intercellular Trafficking. Mol Plant-Microbe Interact (2006) 19:874–883. doi: 10.1094/MPMI-19-0874

19. den Hollander PW, Kieper SN, Borst JW, van Lent Jwm. The role of plasmodesma-located proteins in tubule-guided virus transport is limited to the plasmodesmata. Arch Virol (2016) 161:2431–2440. doi: 10.1007/s00705-016-2936-2

20. den Hollander PW, de Sousa Geraldino Duarte P, Bloksma H, Boeren S, van Lent JWM. Proteomic analysis of the plasma membrane-movement tubule complex of cowpea mosaic virus. Arch Virol (2016) 161:1309–1314. doi: 10.1007/s00705-016-2757-3

21. Fujimoto M, Ueda T. Conserved and Plant-Unique Mechanisms Regulating Plant Post-Golgi Traffic. Front Plant Sci (2012) 3: https://www.frontiersin.org/article/10.3389/fpls.2012.00197

22. Rosquete MR, Drakakaki G. Plant TGN in the stress response: a compartmentalized overview. Curr Opin Plant Biol (2018) 46:122–129. doi: https://doi.org/10.1016/j.pbi.2018.09.003

23. Viotti C, Bubeck J, Stierhof Y-D, Krebs M, Langhans M, van den Berg W, van Dongen W, Richter S, Geldner N, Takano J, et al. Endocytic and Secretory Traffic in Arabidopsis Merge in the Trans-Golgi Network/Early Endosome, an Independent and Highly Dynamic Organelle. Plant Cell (2010) 22:1344–1357. doi: 10.1105/tpc.109.072637

24. Arora D, Damme D Van. Motif-based endomembrane trafficking. Plant Physiol (2021) 186:221–238. doi: 10.1093/plphys/kiab077

25. Geldner N, Robatzek S. Plant Receptors Go Endosomal: A Moving View on Signal Transduction. Plant Physiol (2008) 147:1565–1574. doi: 10.1104/pp.108.120287

26. Junpei T, Mayuki T, Atsushi T, Kyoko M, Koji K, Kentaro F, Hitoshi O, Satoshi N, Toru F. Polar localization and degradation of Arabidopsis boron transporters through distinct trafficking pathways. Proc Natl Acad Sci (2010) 107:5220–5225. doi: 10.1073/pnas.0910744107

27. Gershlick DC, Lousa C de M, Foresti O, Lee AJ, Pereira EA, daSilva LLP, Bottanelli F, Denecke J. Golgi-Dependent Transport of Vacuolar Sorting Receptors Is Regulated by COPII, AP1, and AP4 Protein Complexes in Tobacco. Plant Cell (2014) 26:1308–1329. doi: 10.1105/tpc.113.122226

28. Happel N, Höning S, Neuhaus J-M, Paris N, Robinson DG, Holstein SEH. ArabidopsisµA-adaptin interacts with the tyrosine motif of the vacuolar sorting receptor VSR-PS1. Plant J (2004) 37:678–693. doi: https://doi.org/10.1111/j.1365-313X.2003.01995.x

29. Nishimura K, Matsunami E, Yoshida S, Kohata S, Yamauchi J, Jisaka M, Nagaya T, Yokota K, Nakagawa T. The tyrosine-sorting motif of the vacuolar sorting receptor VSR4 from Arabidopsis thaliana, which is involved in the interaction between VSR4 and AP1M2, μ1-adaptin type 2 of clathrin adaptor complex 1 subunits, participates in the post-Golgi sorting of VS. Biosci Biotechnol Biochem (2016) 80:694–705. doi: 10.1080/09168451.2015.1116925

30. Holstein SEH. Clathrin and Plant Endocytosis. Traffic (2002) 3:614–620. doi: https://doi.org/10.1034/j.1600-0854.2002.30903.x

31. Yoshinari A, Hosokawa T, Amano T, Beier MP, Kunieda T, Shimada T, Hara-Nishimura I, Naito S, Takano J. Polar Localization of the Borate Exporter BOR1 Requires AP2-Dependent Endocytosis. Plant Physiol (2019) 179:1569–1580. doi: 10.1104/pp.18.01017

32. Liu D, Kumar R, Claus LAN, Johnson AJ, Siao W, Vanhoutte I, Wang P, Bender KW, Yperman K, Martins S, et al. Endocytosis of BRASSINOSTEROID INSENSITIVE1 Is Partly Driven by a Canonical Tyr-Based Motif. Plant Cell (2020) 32:3598–3612. doi: 10.1105/tpc.20.00384

33. Solovyev AG, Stroganova TA, Zamyatnin AA, Fedorkin ON, Schiemann J, Morozov SY. Subcellular Sorting of Small Membrane-Associated Triple Gene Block Proteins: TGBp3-Assisted Targeting of TGBp2. Virology (2000) 269:113–127. doi: https://doi.org/10.1006/viro.2000.0200

34. Haupt S, Cowan GH, Ziegler A, Roberts AG, Oparka KJ, Torrance L. Two Plant–Viral Movement Proteins Traffic in the Endocytic Recycling Pathway. Plant Cell (2005) 17:164–181. doi: 10.1105/tpc.104.027821

35. Tilsner J, Cowan GH, Roberts AG, Chapman SN, Ziegler A, Savenkov E, Torrance L. Plasmodesmal targeting and intercellular movement of potato mop-top pomovirus is mediated by a membrane anchored tyrosine-based motif on the lumenal side of the endoplasmic reticulum and the C-terminal transmembrane domain in the TGB3 movement protein. Virology (2010) 402:41–51. doi: https://doi.org/10.1016/j.virol.2010.03.008

36. Rastgou M, Habibi MK, Izadpanah K, Masenga V, Milne RG, Wolf YI, Koonin E V., Turina M. Molecular characterization of the plant virus genus Ourmiavirus and evidence of inter-kingdom reassortment of viral genome segments as its possible route of origin. J Gen Virol (2009) 90:2525–2535. doi: 10.1099/vir.0.013086-0

37. Margaria P, Anderson CT, Turina M, Rosa C. Identification of Ourmiavirus 30K movement protein amino acid residues involved in symptomatology, viral movement, subcellular localization and tubule formation. Mol Plant Pathol (2016) 17:1063–1079. doi: https://doi.org/10.1111/mpp.12348

38. Giulia C, Marina C, Andrea G, Vera M, Massimo T. Reverse Genetic Analysis of Ourmiaviruses Reveals the Nucleolar Localization of the Coat Protein in Nicotiana benthamiana and Unusual Requirements for Virion Formation. J Virol (2011) 85:5091–5104. doi: 10.1128/JVI.02565-10

39. Dettmer J, Hong-Hermesdorf A, Stierhof Y-D, Schumacher K. Vacuolar H+-ATPase Activity Is Required for Endocytic and Secretory Trafficking in Arabidopsis. Plant Cell (2006) 18:715–730. doi: 10.1105/tpc.105.037978

40. Geldner N, Anders N, Wolters H, Keicher J, Kornberger W, Muller P, Delbarre A, Ueda T, Nakano A, Jürgens G. The Arabidopsis GNOM ARF-GEF mediates endosomal recycling, auxin transport, and auxin-dependent plant growth. Cell (2003) 112:219–230. doi: 10.1016/S0092-8674(03)00003-5

41. Jaillais Y, Fobis-Loisy I, Miège C, Rollin C, Gaude T. AtSNX1 defines an endosome for auxin-carrier trafficking in Arabidopsis. Nature (2006) 443:106–109. doi: 10.1038/nature05046

42. Geldner N, Dénervaud-Tendon V, Hyman DL, Mayer U, Stierhof Y-D, Chory J. Rapid, combinatorial analysis of membrane compartments in intact plants with a multicolor marker set. Plant J (2009) 59:169–178. doi: https://doi.org/10.1111/j.1365-313X.2009.03851.x

43. Uemura T, Ueda T, Ohniwa RL, Nakano A, Takeyasu K, Sato MH. Systematic Analysis of SNARE Molecules in Arabidopsis: Dissection of the post-Golgi Network in Plant Cells. Cell Struct Funct (2004) 29:49–65. doi: 10.1247/csf.29.49

44. Sohn EJ, Kim ES, Zhao M, Kim SJ, Kim H, Kim Y-W, Lee YJ, Hillmer S, Sohn U, Jiang L, et al. Rha1, an Arabidopsis Rab5 Homolog, Plays a Critical Role in the Vacuolar Trafficking of Soluble Cargo Proteins. Plant Cell (2003) 15:1057–1070. doi: 10.1105/tpc.009779

45. Ito Y, Toyooka K, Fujimoto M, Ueda T, Uemura T, Nakano A. The trans-Golgi Network and the Golgi Stacks Behave Independently During Regeneration After Brefeldin A Treatment in Tobacco BY-2 Cells. Plant Cell Physiol (2017) 58:811–821. doi: 10.1093/pcp/pcx028

46. Wang J, Cai Y, Miao Y, Lam SK, Jiang L. Wortmannin induces homotypic fusion of plant prevacuolar compartments*. J Exp Bot (2009) 60:3075–3083. doi: 10.1093/jxb/erp136

47. Zheng J, Han SW, Rodriguez-Welsh MF, Rojas-Pierce M. Homotypic Vacuole Fusion Requires VTI11 and Is Regulated by Phosphoinositides. Mol Plant (2014) 7:1026–1040. doi: https://doi.org/10.1093/mp/ssu019

48. Nelson BK, Cai X, Nebenführ A. A multicolored set of in vivo organelle markers for co-localization studies in Arabidopsis and other plants. Plant J (2007) 51:1126–1136. doi: https://doi.org/10.1111/j.1365-313X.2007.03212.x

49. Robinson DG, Langhans M, Saint-Jore-Dupas C, Hawes C. BFA effects are tissue and not just plant specific. Trends Plant Sci (2008) 13:405–408. doi: 10.1016/j.tplants.2008.05.010

50. Langhans M, Förster S, Helmchen G, Robinson DG. Differential effects of the brefeldin A analogue (6R)-hydroxy-BFA in tobacco and Arabidopsis. J Exp Bot (2011) 62:2949–2957. doi: 10.1093/jxb/err007

51. Pouwels J, Van Der Krogt GNM, Van Lent J, Bisseling T, Wellink J. The Cytoskeleton and the Secretory Pathway Are Not Involved in Targeting the Cowpea Mosaic Virus Movement Protein to the Cell Periphery. Virology (2002) 297:48–56. doi: https://doi.org/10.1006/viro.2002.1424

52. Bonifacino JS, Traub LM. Signals for Sorting of Transmembrane Proteins to Endosomes and Lysosomes. Annu Rev Biochem (2003) 72:395–447. doi: 10.1146/annurev.biochem.72.121801.161800

53. Law KC, Chung KK, Zhuang X. An Update on Coat Protein Complexes for Vesicle Formation in Plant Post-Golgi Trafficking. Front Plant Sci (2022) 13:826007. doi: 10.3389/fpls.2022.826007

54. Di Rubbo S, Irani NG, Kim SY, Xu Z-Y, Gadeyne A, Dejonghe W, Vanhoutte I, Persiau G, Eeckhout D, Simon S, et al. The Clathrin Adaptor Complex AP-2 Mediates Endocytosis of BRASSINOSTEROID INSENSITIVE1 in Arabidopsis. Plant Cell (2013) 25:2986–2997. doi: 10.1105/tpc.113.114058

55. Fan L, Hao H, Xue Y, Zhang L, Song K, Ding Z, Botella MA, Wang H, Lin J. Dynamic analysis of Arabidopsis AP2 σ subunit reveals a key role in clathrin-mediated endocytosis and plant development. Development (2013) 140:3826–3837. doi: 10.1242/dev.095711

56. Kim SY, Xu Z-Y, Song K, Kim DH, Kang H, Reichardt I, Sohn EJ, Friml J, Juergens G, Hwang I. Adaptor Protein Complex 2–Mediated Endocytosis Is Crucial for Male Reproductive Organ Development in Arabidopsis. Plant Cell (2013) 25:2970–2985. doi: 10.1105/tpc.113.114264

57. Yamaoka S, Shimono Y, Shirakawa M, Fukao Y, Kawase T, Hatsugai N, Tamura K, Shimada T, Hara-Nishimura I. Identification and Dynamics of Arabidopsis Adaptor Protein-2 Complex and Its Involvement in Floral Organ Development. Plant Cell (2013) 25:2958–2969. doi: 10.1105/tpc.113.114082

58. Gadeyne A, Sánchez-Rodríguez C, Vanneste S, Di Rubbo S, Zauber H, Vanneste K, Van Leene J, De Winne N, Eeckhout D, Persiau G, et al. The TPLATE Adaptor Complex Drives Clathrin-Mediated Endocytosis in Plants. Cell (2014) 156:691–704. doi: 10.1016/j.cell.2014.01.039

59. Wang C, Hu T, Yan X, Meng T, Wang Y, Wang Q, Zhang X, Gu Y, Sánchez-Rodríguez C, Gadeyne A, et al. Differential Regulation of Clathrin and Its Adaptor Proteins during Membrane Recruitment for Endocytosis. Plant Physiol (2016) 171:215–229. doi: 10.1104/pp.15.01716

60. Shemyakina EA, Erokhina TN, Gorshkova EN, Schiemann J, Solovyev AG, Morozov SY. Formation of protein complexes containing plant virus movement protein TGBp3 is necessary for its intracellular trafficking. Biochimie (2011) 93:742–748. doi: https://doi.org/10.1016/j.biochi.2011.01.002

61. Wang X, Cai Y, Wang H, Zeng Y, Zhuang X, Li B, Jiang L. Trans-Golgi Network-Located AP1 Gamma Adaptins Mediate Dileucine Motif-Directed Vacuolar Targeting in Arabidopsis. Plant Cell (2014) 26:4102–4118. doi: 10.1105/tpc.114.129759

62. Müdsam C, Wollschläger P, Sauer N, Schneider S. Sorting of Arabidopsis NRAMP3 and NRAMP4 depends on adaptor protein complex AP4 and a dileucine-based motif. Traffic (2018) 19:503–521.

63. Wu G, Cui X, Dai Z, He R, Li Y, Yu K, Bernards M, Chen X, Wang A. A plant RNA virus hijacks endocytic proteins to establish its infection in plants. Plant J (2020) 101:384–400. doi: https://doi.org/10.1111/tpj.14549

64. Wu G, Xiaoyan C, Chen H, Renaud JB, Yu K, Chen X, Wang A. Dynamin-Like Proteins of Endocytosis in Plants Are Coopted by Potyviruses To Enhance Virus Infection. J Virol (2018) 92:e01320–18. doi: 10.1128/JVI.01320-18

65. Levy A, Tilsner J. Creating Contacts Between Replication and Movement at Plasmodesmata - A Role for Membrane Contact Sites in Plant Virus Infections? Front Plant Sci (2020) 11:862. doi: 10.3389/fpls.2020.00862

66. Cheng Y, G. Ls, Vitaly C, E. Ls. Identification of a Functional Plasmodesmal Localization Signal in a Plant Viral Cell-To-Cell-Movement Protein. MBio (2016) 7:e02052–15. doi: 10.1128/mBio.02052-15

67. Bashline L, Li S, Anderson CT, Lei L, Gu Y. The Endocytosis of Cellulose Synthase in Arabidopsis Is Dependent on μ2, a Clathrin-Mediated Endocytosis Adaptin. Plant Physiol (2013) 163:150–160. doi: 10.1104/pp.113.221234

68. Coutu C, Brandle J, Brown D, Brown K, Miki B, Simmonds J, Hegedus DD. pORE: a modular binary vector series suited for both monocot and dicot plant transformation. Transgenic Res (2007) 16:771–781. doi: 10.1007/s11248-007-9066-2

