## Supplementary for "Sorting motifs target the movement protein of ourmia melon virus to the trans-Golgi network and plasmodesmata"

### Supplementary figure legends

**Figure S1. Protein lipid overlay assay demonstrates the binding of MBP-MP to PI3P.** LPA: Lysophosphatidic Acid, LPC: Lysophosphocholine, PI: Phosphatidylinositol, PI3P: Phosphatidylinositol 3-phosphate, PI4P: Phosphatidylinositol 4-phosphate, PI5P: Phosphatidylinositol 5-phosphate, PE: Phosphatidylethanolamine, PC: Phosphatidylcholine, S1P: Sphingosine-1-phosphate, PI(3,4)P2: Phosphatidylinositol 3,4-bisphosphate, PI(3,5)P2: Phosphatidylinositol 3,5-bisphosphate, PI(4,5)P2: Phosphatidylinositol 4,5-bisphosphate, PI(3,4,5)P3: Phosphatidylinositol 3,4,5-trisphosphate, PA: Phosphatidic Acid, PS: Phosphatidylserine.

**Figure S2. Symptoms of *N. benthamiana* plants infected with OuMV carrying MP_D/G. (A)** Magnified image of the leaf in Figure 2C. **(B)** Viral symptoms appeared at 5 dpi. White arrows show upper uninoculated leaves and white plus signs indicate agroinfiltrated leaves. The background was removed from the images.

**Figure S3. Agroinfiltrated areas are saturated with GFP.** Representative images of *N. benthamiana* epidermal cells expressing pGC-RNA2 (MP) or mutants, together with pGC-RNA1 (RdRP), pGC-RNA3 (CP) and pBin61-GFP. Images represent single optical planes. Scale bars, 50 μm.

**Figure S4. Replication assay of OuMV carrying Y and LL motif MP mutants at 24 hpi.** Two leaf spots from 3 leaves were pooled for each sample at 24 hpi and 48 hpi. The qPCR was performed in duplicates. The $2^{-\Delta\Delta Ct}$ method was used to assess fold changes in replication of wild-type and mutants. The data was normalized first using 18S ribosomal RNA, and then a wild-type control at 24 hpi showing the lowest expression. Box plot represents the relative expression of RNA1 (n=4). The statistical analysis was performed on ΔΔCt values using Student's t-test (unpaired, two-tailed).

**Figure S5. Time-dependent intracellular localization of MP mutants fused to GFP in *N. benthamiana* epidermal cells.** Fluorescent patterns were monitored every 12 h starting from 36 h to 72 h following agroinfiltration of bacterial cultures carrying plasmids pGC-RNA1(RdRP), pGC-RNA3 (CP) and pGC-RNA2 expressing GFP:MPwt **(A)** or GFP:MP mutants, Y (**B-C)** and LL **(D-F)** sorting motif mutants. Images represent maximum intensity Z projections. White arrowheads denote puncta. Scale bars, 10 μm.

**Figure S6. Y motif mutants are not targeted to the plasma membrane.** *N.benthamiana* epidermal cells expressing **(A)** GFP:MPwt or GFP:MP mutants **(B)** GFP:MP_Y and **(C)** GFP:MP_Y/G, along with RdRP and CP. Cells were stained with FM4-64, a plasma membrane marker (magenta). The intensity profile of the diagonal line was generated by ImageJ. Images were taken at 66 hpi and represent single optical planes. Scale bar, 10 μm.

**Movies**

**Movie S1. Time lapse imaging of GFP:MPwt in *N. benthamiana* epidermal cells showing mobile punctate structures at the cell periphery.** The images were taken at 60 hpi and were acquired at intervals of 1 s over a period of 15 s. Scale bar, 10 μm.

**Movie S2. Time lapse imaging of GFP:MPwt in *N. benthamiana* epidermal cells showing cytoplasmic mobile punctate structures**. The images were taken at 60hpi and were acquired at intervals of 1 s over a period of 15 s. Scale bar, 10 μm.

**Movie S3. Time lapse imaging of Y motif mutants in *N. benthamiana* epidermal cells showing the cytoplasmic localization and absence of punctate structures.** Occasionally, GFP:MP_Y/G labels stationary or less mobile puncta. The images were taken at 60 hpi and were acquired at intervals of 1 s over a period of 15 s. Scale bar, 10 μm.

**Movie S4. Time lapse imaging of LL motif mutants in *N. benthamiana* epidermal cells showing the presence of mobile puncta**. The images were taken at 60 hpi and were acquired at intervals of 1 s over a period of 15 s. Scale bar, 10 μm.
