## Supplementary figures and images for "Sorting motifs target the movement protein of ourmia melon virus to the trans-Golgi network and plasmodesmata"

### Fig. S1

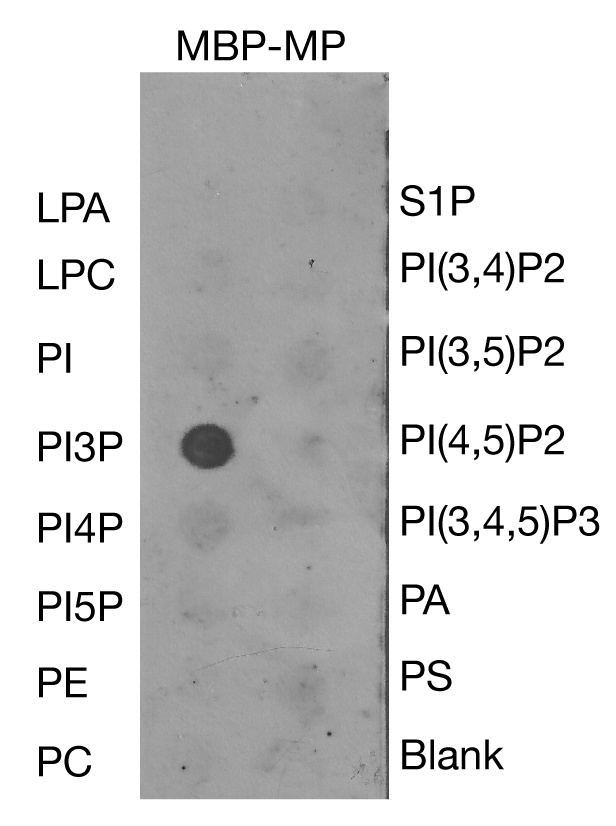

### Fig. S2

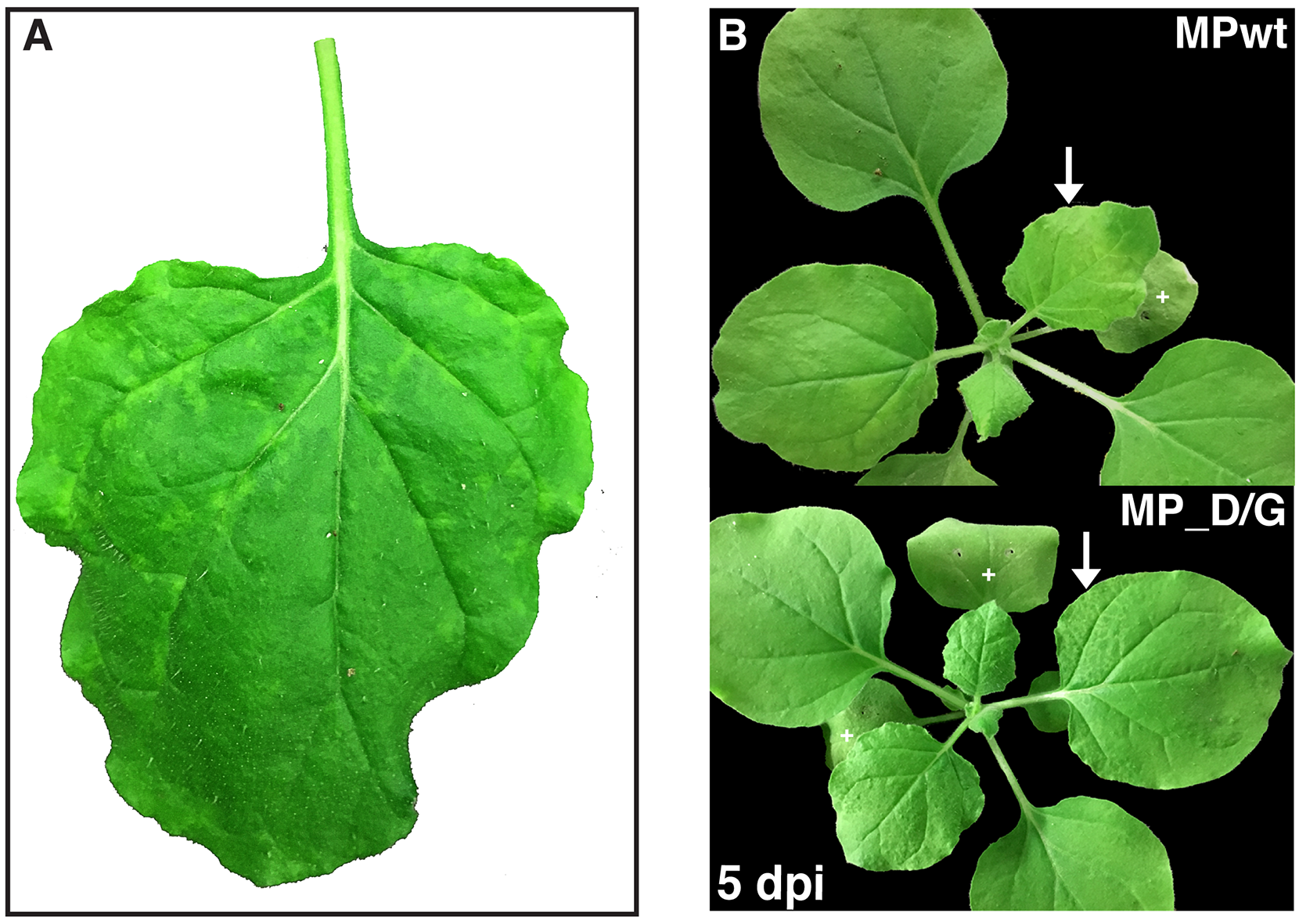

### Fig. S3

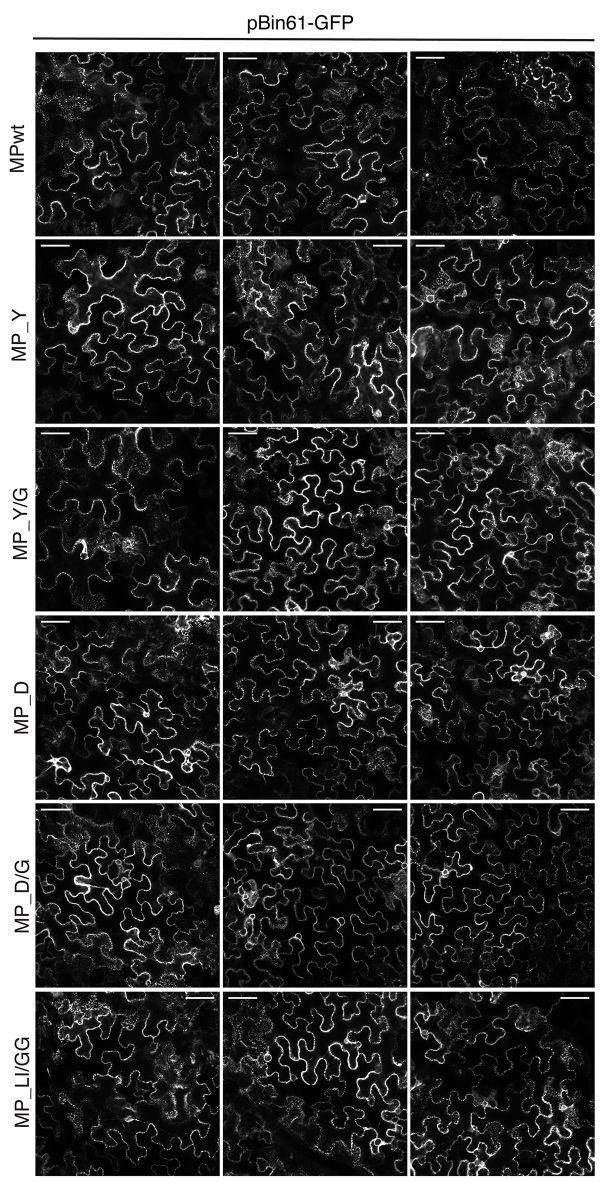

### Fig. S4

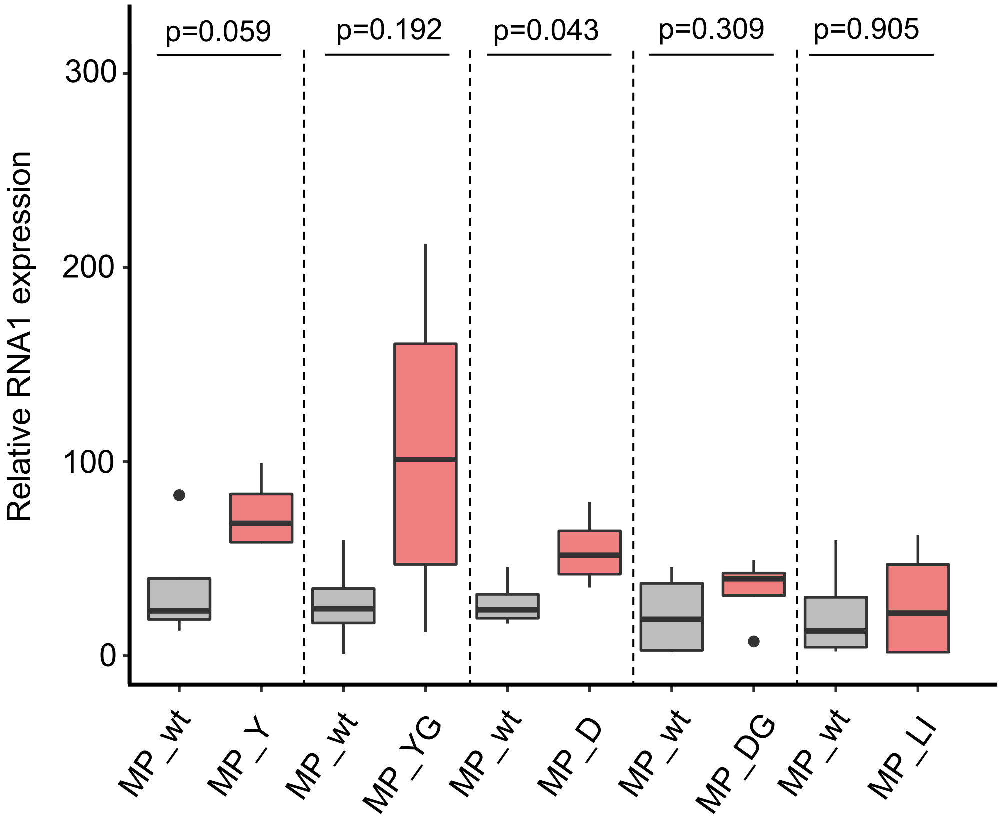

### Fig. S5

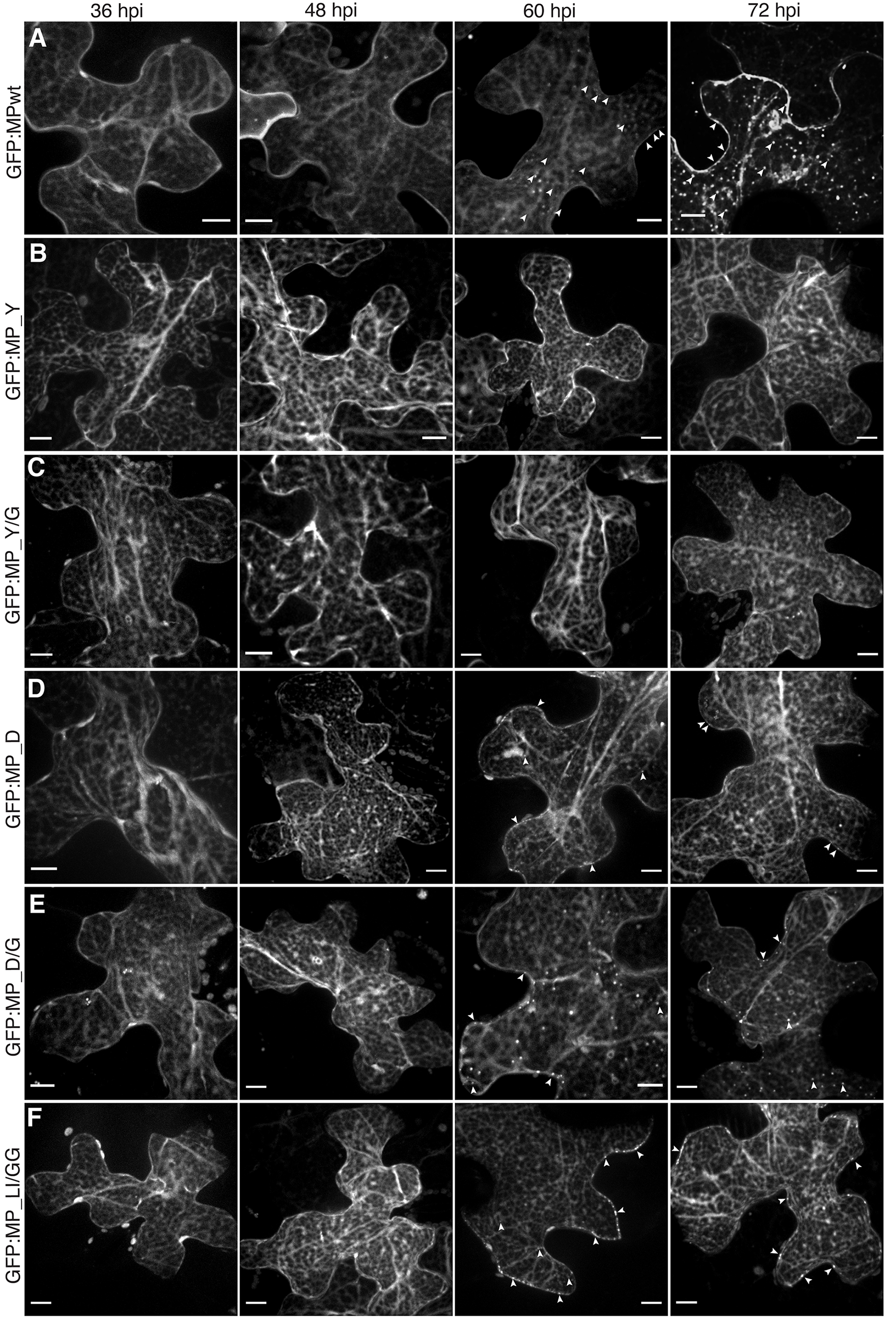

### Fig. S6

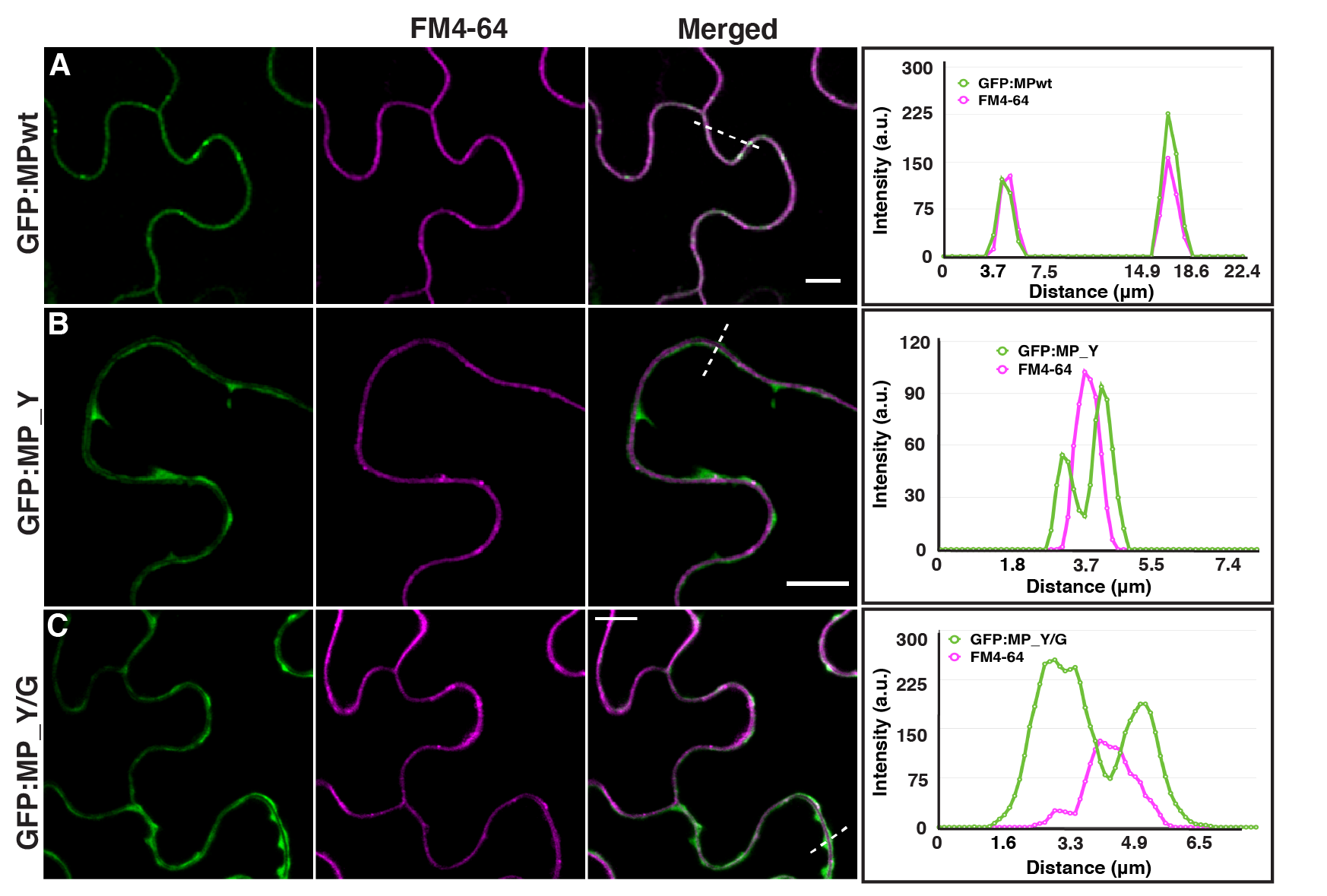
