## Supplementary material for "Sorting motifs target the movement protein of ourmia melon virus to the trans-Golgi network and plasmodesmata": Table S1

Table 1. Candidate detergent-soluble OuMV MP-interacting proteins^a,b^

| Function | Gene locus | Protein name | Occurrence/Total replicates |
| --- | --- | --- | --- |
| Vesicular trafficking |  |  |  |
|  | AT3G54840 | ARA6 | 1/3 |
| Ca^2+^ signaling |  |  |  |
|  | ^c^AT5G37780 AT2G41110  AT3G56800  AT1G66410  AT2G27030  AT5G21274  AT3G43810 | Calmodulin  (CAM 1-7) | 1/3 |
| Transport |  |  |  |
|  | ^c^AT2G18960  AT4G30190  AT5G57350 AT3G47950  AT3G42640  AT1G17260  AT5G62670 | Plasma membrane proton ATPase  (AHA 1-4,8,10-11) | 1/3 |
|  | AT4G35100 | Plasma membrane intrinsic protein 3A (PIP3A, PIP2;7) | 1/3 |
|  | ^c^AT1G76030  AT4G38510  AT1G20260 | V-type proton ATPase subunit B  (VAB1/2/3) | 1/3 |
| Chaperones |  |  |  |
|  | ^c^AT5G2206  AT3G44110 | DNAJ homologue (ATJ2/3) | 3/3 |
|  | ^c^AT5G42020  AT5G28540 | Luminal binding protein (BiP1/2) | 1/3 |
| Metabolic enzymes |  |  |  |
|  | ^c^AT3G14420  AT3G14415  AT4G18360 | (S)-2-hydroxy-acid oxidase  (GLO1/2/3) | 1/3 |
|  | ^c^AT1G02500  AT4G01850 | S- adenosylmethionine synthase (SAM1/2) | 1/3 |
| Redox |  |  |  |
|  | AT3G17880 | Tetraticopeptide domain-containing thioredoxin  (AtTDX) | 3/3 |
| Proteolysis |  |  |  |
|  | AT3G14067 | Senescence-associated subtilisin protease (SASP) | 2/3 |
|  | ^c^AT5G60360  AT3G45310 | Senescence associated gene 2 (SAG2)/ Cysteine proteinases superfamily protein | 1/3 |

a, unused score > 1.3; b, ribosomal proteins, elongation factors, and chloroplast localized proteins were not included; c, identified peptides map to multiple proteins in the same family
