## Supplementary material for "Sorting motifs target the movement protein of ourmia melon virus to the trans-Golgi network and plasmodesmata": Table S2

Table 2. Candidate soluble OuMV MP-interacting proteins^a,b^

| Function | Gene  locus | Protein  name | Occurrence /Total replicates |
| --- | --- | --- | --- |
| Vesicular trafficking |  |  |  |
|  | AT1G60070  AT5G22780  AT1G31730 | AP1, gamma subunit 2  AP2, alpha subunit  AP4, epsilon | 1/3  1/3  1/3 |
|  | AT5G41950  AT1G72150  AT2G27600  ^c^AT4G14160  AT1G05520  ^c^AT4G17890  AT5G46750  AT5G51430 | Hypersensitivity to latrunculin B1 (HLB1)  Patellin 1  (PATL1)  Suppressor of K+ transport growth defect 1  (SKD1)  Sec23/Sec24  Probable ADP-ribosylation factor GTPase-activating protein (AGD8/9)  EYE | 1/3  1/3  1/3  1/3  1/3  1/3 |
| Lipid binding |  |  |  |
|  | AT3G19420  AT5G07370    AT5G61760 | Phosphatidylinositol 3,4,5-trisphosphate 3-phosphatase and protein-tyrosine-phosphatase (PTEN2A)  Inositol polyphosphate 6-/3-/5-kinase alpha  (IPK2α)  Inositol polyphosphate 6-/3-/5-kinase (IPK2β) | 1/3  2/3  2/3 |
| Kinase/  Phosphatase |  |  |  |
|  | AT1G71860 | Protein tyrosine phosphatase 1 | 1/3 |
|  | AT2G42500 | Protein phosphatase 2A-3 | 1/3 |
|  | AT1G51690  AT1G30470  AT1G60940 | Serine/threonine-protein phosphatase 2A 55 kDa regulatory subunit B  SIT4 phosphatase- associated family protein  Serine/threonine-protein kinase SRK2B  (SNRK 2.10) | 1/3  1/3  1/3 |
| Chaperones |  |  |  |
|  | AT3G09440 | Heat shock 70 kDa protein 3 | 2/3 |
|  | ^c^AT5G56000  AT5G56010  AT5G56030 | Heat shock protein 81  2-4 | 1/3 |
| RNA binding |  |  |  |
|  | AT5G61780  AT3G04610 | TUDOR-SN protein 2  RNA-binding KH domain-containing (FLK) | 1/3  2/3 |
| Cell wall  related |  |  |  |
|  | AT2G06850  AT1G72680  AT1G15950 | Xyloglucan endotransglucosylase/hydrolase 4  (XHT4)  Cinnamyl-alcohol dehydrogenase  (CAD 1)  Cinnamoyl-CoA reductase 1 (CCR1/IRX4) | 1/3  2/3  1/3 |
| Protein degradation |  |  |  |
|  | ^c^AT3G52590  AT2G36170 | Ubiquitin extension protein 1/2  (UBQ1/2) | 1/3 |
|  | ^c^AT1G55860  AT1G70320 | Ubiquitin-protein ligase 1/2  (UPL1/2) | 2/3 |
|  | AT1G70320 | Ubiquitin-protein ligase 2  (UPL2) | 1/3 |
|  | AT4G28470  AT5G09900  AT1G28120  ^c^AT3G51260  AT5G66140  AT3G09840  AT2G39940 | 26S proteasome non-ATPase regulatory subunit 2 homolog B  (RPN1B)  26S proteasome non-ATPase regulatory subunit 12 homolog A  (RPN5A)  Ubiquitin thioesterase otubain-like protein  Proteasome subunit alpha type  (PAD1/2)  Cell division control protein 48  (CDC48)  Coronatine-insensitive protein 1  (COI1) | 1/3  2/3  3/3  1/3  1/3  1/3 |
| Blue light  response |  |  |  |
|  | AT3G45780  AT3G07640 | Phototropin 1  (PHOT1)  Period circadian  protein | 1/3  1/3 |

a, unused score > 1.3; b, ribosomal proteins, elongation factors, and chloroplast localized proteins were not included; c, identified peptides map to multiple proteins in the same family
