## Supplementary material for "Sorting motifs target the movement protein of ourmia melon virus to the trans-Golgi network and plasmodesmata": Table S3

**Table S3.** Systemic infection by OuMV in Arabidopsis Col-0 (wt), *ap2m-1* mutant and AP2M-YFP complementation line at 14 dpi.

|  | Systemically infected plants/ total inoculated plants |
| --- | --- |
| Col-0 (wt) | 7/12 |
| *ap2m-1* | 7/12 |
| AP2M-YFP/*ap2m-1* | 6/12 |
