## Supplementary material for "Sorting motifs target the movement protein of ourmia melon virus to the trans-Golgi network and plasmodesmata": Table S4

**Table S4. Subcellular localization of Y and LL motif mutants**

|  | Cytoplasmic Punctae | PM | PD |
| --- | --- | --- | --- |
| OuMV-MPwt | **+**  **(TGN/endosomes and occasional immobile)** | **+** | **+** |
| OuMV-MP_Y | **-** | **-** | **-** |
| OuMV-MP_ Y/G | **-**  **(a few, immobile)** | **-** | **-** |
| OuMV-MP_D | **+**  **(mobile, occasional immobile)** | **+** | **-** |
| OuMV-MP_D/G | **+**  **(mobile, occasional immobile)** | **+** | **+** |
| OuMV-MP_LI/GG | **+**  **(mobile, occasional immobile)** | **+** | **-** |
