## Supplementary material for "Sorting motifs target the movement protein of ourmia melon virus to the trans-Golgi network and plasmodesmata": Table S5

**Table S5. List of primers used in this study.**

| Primer | Sequence (5'-3') |  |
| --- | --- | --- |
| Y | F: AAGGGAATTCTCTCTTGGAAACCG  R: ATCACCGGTAATGTATCCTTTTGGG | To insert the mutations Y88G and V91G |
| Y/G | F: ATACATTACCGGTGATAAGGTAATTCTCTCTTGGAAACC  R: CCTTTTGGGCATCGCAGT | To insert the mutation Y88G |
| D | F: GCCGGGGGTCCGCACGGTATCTGGTCC  R: GATCGGGCCCTCGACATTAGCATCTAGCGAAATG | To insert the mutations D59G and LI63-64GG |
| D/G | F: TAATGTCGAGGGCCCGATCGCC  R: GCATCTAGCGAAATGTCGTGG | To insert the mutation D59G |
| LI/GG | F: CCCGATCGCCGGGGGTCCGCACGGTATC  R: TCCTCGACATTAGCATCTAG | To insert the mutations LI63-64GG |
| Ubq10  XhoI | F: AGCTACTCGAGAGTCTAGCTCAACAGAGCTTTTAAC | To insert Ubq10pro::mCherry::VTI12 and Ubq10pro::mCherry::RHA1 into pORE O2 |
| VTI12  SpeI | R: TAGCTACTAGTTTAATGAGAAAGCTTGTATGAGATG |  |
| RHA1  SpeI | R: TAGCTACTAGTCTAAGCACAACACGATGAACTCA |  |
| MP | F: ATGGGGGACAATGCTTTAGA  R: CCTGACCGAAGCCAGGAGAA | To amplify full-length MP from cDNAs of MP mutants |
| MP84 | F: TAAGAGATGTAAGCCAATCCTG  R: TCCTTTTGGGCATCGCAG | To delete 85-288 region of MP |
| MP102 | F: TAAGAGATGTAAGCCAATCCTGTAAAATCCTGG  R: TCCGGTGGCCACATGCGG | To delete 103-288 region of MP |
| Ap2m | LP: GAATCTGGTCAGGTTCACACACTGGTG  RP: GCAATGCTAATGTTGCTTGTG | Genotyping of *ap2m-1* plants |
| RNA1 | F: CGAAGAACTGGGACGAGAAG  R: ATGTCGGATTCTTCCACAGG | To quantify RNA1 for replication assay |
| 18S | F: ATGGCCGTTCTTAGTTGGTGGAGC  R: AGTTAGCAGGCTGAGGTCTCGAAC | Internal control for qPCR |
